# A novel antibody targeting ICOS increases intratumoural cytotoxic to regulatory T cell ratio and induces tumour regression

**DOI:** 10.1101/771493

**Authors:** Richard C.A. Sainson, Anil K. Thotakura, Miha Kosmac, Gwenoline Borhis, Nahida Parveen, Rachael Kimber, Joana Carvalho, Simon Henderson, Kerstin Pryke, Tracey Okell, Siobhan O’Leary, Stuart Ball, Lauriane Gamand, Emma Taggart, Eleanor Pring, Hanif Ali, Hannah Craig, Vivian W. Y. Wong, Qi Liang, Robert J. Rowlands, Morgane Lecointre, Jamie Campbell, Ian Kirby, David Melvin, Volker Germaschewski, Elisabeth Oelmann, Sonia Quaratino, Matthew McCourt

## Abstract

The immunosuppressive tumour microenvironment constitutes a significant hurdle to the response to immune checkpoint inhibitors. Both soluble factors and specialised immune cells such as regulatory T cells (T_Reg_) are key components of active intratumoural immunosuppression. Previous studies have shown that Inducible Co-Stimulatory receptor (ICOS) is highly expressed in the tumour microenvironment, especially on T_Reg_, suggesting that it represents a relevant target for preferential depletion of these cells. Here, we used immune profiling of samples from tumour bearing mice and cancer patients to characterise the expression of ICOS in different tissues and solid tumours. By immunizing an *Icos* knockout transgenic mouse line expressing antibodies with human variable domains, we selected a fully human IgG1 antibody called KY1044 that binds ICOS from different species. Using KY1044, we demonstrated that we can exploit the differential expression of ICOS on T cell subtypes to modify the tumour microenvironment and thereby improve the anti-tumour immune response. We showed that KY1044 induces sustained depletion of ICOS^high^ T_Reg_ cells in mouse tumours and depletion of ICOS^high^ T cells in the blood of non-human primates, but was also associated with secretion of pro-inflammatory cytokines from ICOS^low^ T_EFF_ cells. Altogether, KY1044 improved the intratumoural T_EFF_:T_Reg_ ratio and increased activation of T_EFF_ cells, resulting in monotherapy efficacy or in synergistic combinatorial efficacy when administered with the immune checkpoint blocker anti-PD-L1. In summary, our data demonstrate that targeting ICOS with KY1044 can favourably alter the intratumoural immune contexture, promoting an anti-tumour response.

## Introduction

The last decade has seen a paradigm shift in cancer therapies, with the approval of antibodies targeting immune checkpoints (e.g. CTLA-4, PD-1 and PD-L1). These immune checkpoint inhibitors (ICIs) are associated with strong and durable responses in patients suffering from advanced malignancies, including but not limited to metastatic melanoma, non-small cell lung cancer (NSCLC), head and neck cancer, renal and bladder cancer (*1*). However, within all these indications, there are still a high proportion of patients who exhibit intrinsic or acquired resistance to ICIs. These patients represent a population with high unmet medical needs who may benefit from novel combinatory approaches with ICIs.

Multiple molecular and cellular mechanisms have been associated with the lack of response to immunotherapies (*2*). Accumulating evidence has shown that a low incidence of cytotoxic T cells and the presence of immunosuppressive cells represent major barriers to establishing a response to ICIs. One such class of immunosuppressive cells are regulatory T cells (T_Reg_), which are characterised by the expression of the transcription factor FOXP3 and function by blocking the activation and cytotoxic potential of effector T-cells (T_Eff_) through multiple mechanisms (*3, 4*). In fact, numbers of intratumoural T_Reg_ cells negatively correlate with patient survival in several cancer types including melanoma, NSCLC, breast cancer and hepatocellular carcinoma, where a high T_Reg_ content is associated with poor prognosis and minimal response to standard of care therapy (*5, 6*). Of relevance, T_Reg_ depletion modifies the immune contexture in the tumour microenvironment (TME) and favours an anti-tumour response in several different mouse tumour models (*7–9*). For these reasons, T_Reg_ cells have been investigated as a potential prognostic cell type and as therapeutic targets for depletion strategies.

The use of therapeutic antibodies for the preferential depletion of intratumoral T_Reg_ cells relies on the identification of a marker that is preferably highly expressed on the surface of these cells in the TME compared with other tissues. One such potential target is the Inducible Co-Stimulatory receptor (ICOS) which belongs to the CD28 and CTLA-4 family (*10*). Unlike CD28, ICOS is not expressed on naïve T_Eff_ cells but is rapidly induced upon T cell receptor (TCR) engagement (*11, 12*). Following activation, ICOS has been shown to be constitutively expressed on the surface of T cells where it can engage with its unique ligand (ICOS-LG, CD275) expressed primarily on antigen presenting cells. ICOS/ICOS-LG interaction initiates a co-stimulatory signal that results in production of either pro- or anti-inflammatory cytokines (IFN-*γ* and TNF-α by T_Eff_ cells; IL-10 expression by T_Reg_) (*13*). Overall the ICOS/ICOS-LG signalling pathway has been shown to regulate immune cell homeostasis and be implicated in the overall immune response (*14, 15*). Of relevance, the accumulation of ICOS^+^ T_Reg_ cells in the TME has been shown to be associated with disease progression in cancer patients (*16, 17*). In marked contrast, the upregulation of ICOS on CD4 T_EFF_ cells has been associated with an anti-tumour response and better prognosis in patients after treatment with anti-CTLA-4 (*18–21*). Interestingly, the relative expression levels of ICOS vary between different T cell subtypes, with T_Reg_ exhibiting the highest levels of ICOS followed by CD4^+^ and then CD8^+^ T_EFF_ cells (*19*). Similarly, some reports have suggested that ICOS expression is higher in the TME than in other tissues (*12, 17, 22*). This differential expression between T cell subtypes, and the high expression levels of ICOS in intratumoural T_Reg_ cells (*22*), suggests that ICOS represents a relevant target for a T_Reg_ depletion strategy. In fact, it has been demonstrated that anti-ICOS antibodies with depleting capability can specifically reduce numbers of ICOS^+^ T_Reg_ cells and induce a strong anti-tumour immune response when combined with a vaccine strategy (*7*). Given the co-stimulatory and pro-inflammatory effects of ICOS signalling in T_EFF_ cells, an ideal anti-cancer therapeutic strategy would be to promote the depletion of ICOS^High^ T_Reg_ cells while boosting the proinflammatory potential of ICOS^Low^ CD8 T_EFF_ cells in the TME.

In the present study, we used our antibody generating transgenic mouse platform (*23, 24*) to identify a fully human anti-ICOS IgG1 antibody called KY1044 that binds with similar affinity to mouse and human ICOS. *In vitro*, we established that KY1044 has a dual mechanism of action, namely a co-stimulatory effect on ICOS^Low^ T_EFF_ cells and a depleting effect on ICOS^High^ T_Reg_. *In vivo,* pharmacodynamic analysis confirmed that KY1044 induces preferential T_Reg_ depletion and increased secretion of pro-inflammatory cytokines by activated T_EFF_ cells in the TME. Overall, KY1044 was shown to modulate the TME immune contexture by increasing the CD8^+^:T_Reg_ cell ratio. Importantly, this change in ratio was associated with anti-tumour efficacy when KY1044 was used as monotherapy and with synergistic activity when KY1044 was combined with anti-PD-L1. Altogether, our results highlight the potential of KY1044 as an anti-tumour therapeutic agent to diminish immune suppression in the TME and re-establish anti-tumour immunity in patients.

## Results

### ICOS is highly expressed on intratumoural T_Reg_ in several cancer indications

Many tumour types, such as ovarian, gastric and liver cancers, contain a high number of ICOS^+^ T_Reg_ cells (*16, 17, 25*). A high number of these cells in the TME is associated with poor prognosis, possibly due to the fact that they are highly immunosuppressive (*17, 26*). In order to assess ICOS expression on intratumoural T cells and to identify additional tumour types containing high ICOS^+^ T_Reg_ cells, we first analysed the content of T cells in tumours using different approaches.

We initially interrogated the TCGA (The Cancer Genome Atlas) mRNA datasets to identify tumours with high levels of *Icos* and *Foxp3* (a marker of T_Reg_) mRNAs (Fig. 1A). Several cancers expressed high levels of these transcripts including tumours originating from the head and neck, stomach, cervix, thymus, testis, skin and lung. Since this approach does not establish which types of cells from the tumour expressed ICOS and/or FOXP3, we further assessed these markers at the mRNA and protein levels by single-cell transcriptomics and FACS analysis respectively. We obtained paired peripheral blood mononuclear cell (PBMC) and tumour scRNA-seq data for 79,544 cells from 5 NSCLC patients, which we further stratified into 27 immune cell subtypes based on their gene expression signatures (Supplementary Table 1). As expected, certain genes such as *foxp3*, *ccr8*, *il2ra*, *tnfrsf4* were shown to be highly expressed in intra-tumoural T_Reg_ thus validating the approach used (Fig. 1B). *Icos* mRNA expression in these NSCLC samples was higher in T_Regs_ than in other T cell subsets. Finally, *Icos* gene expression levels were also higher in the TME than in the periphery while the pattern of expression in both compartments followed the general trend of: T_Reg_ > CD4_non-TReg_ > CD8 > Other (Fig. 1B) suggesting that one could exploit the differential expression between the T cell subsets.

**Figure 1:**
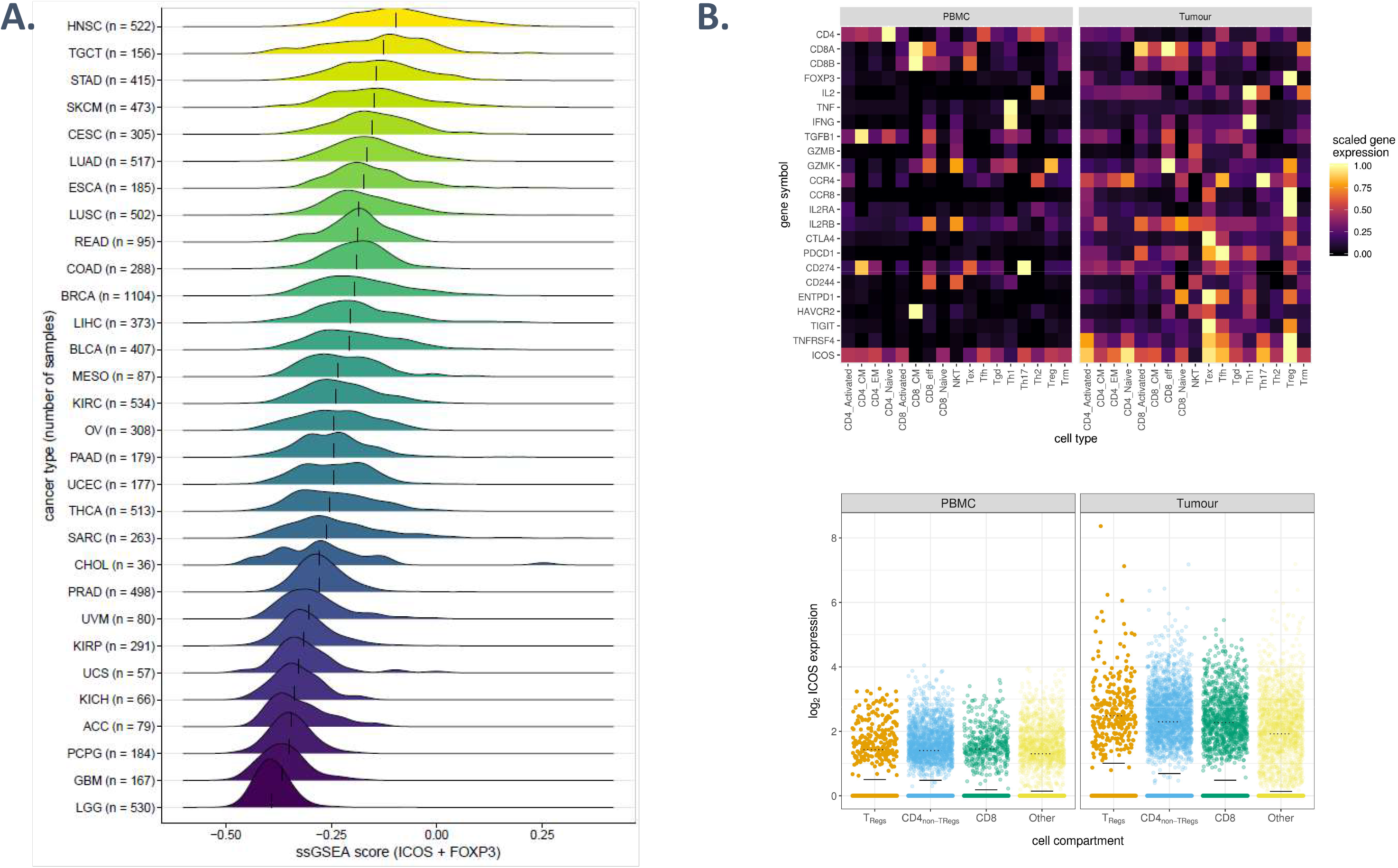
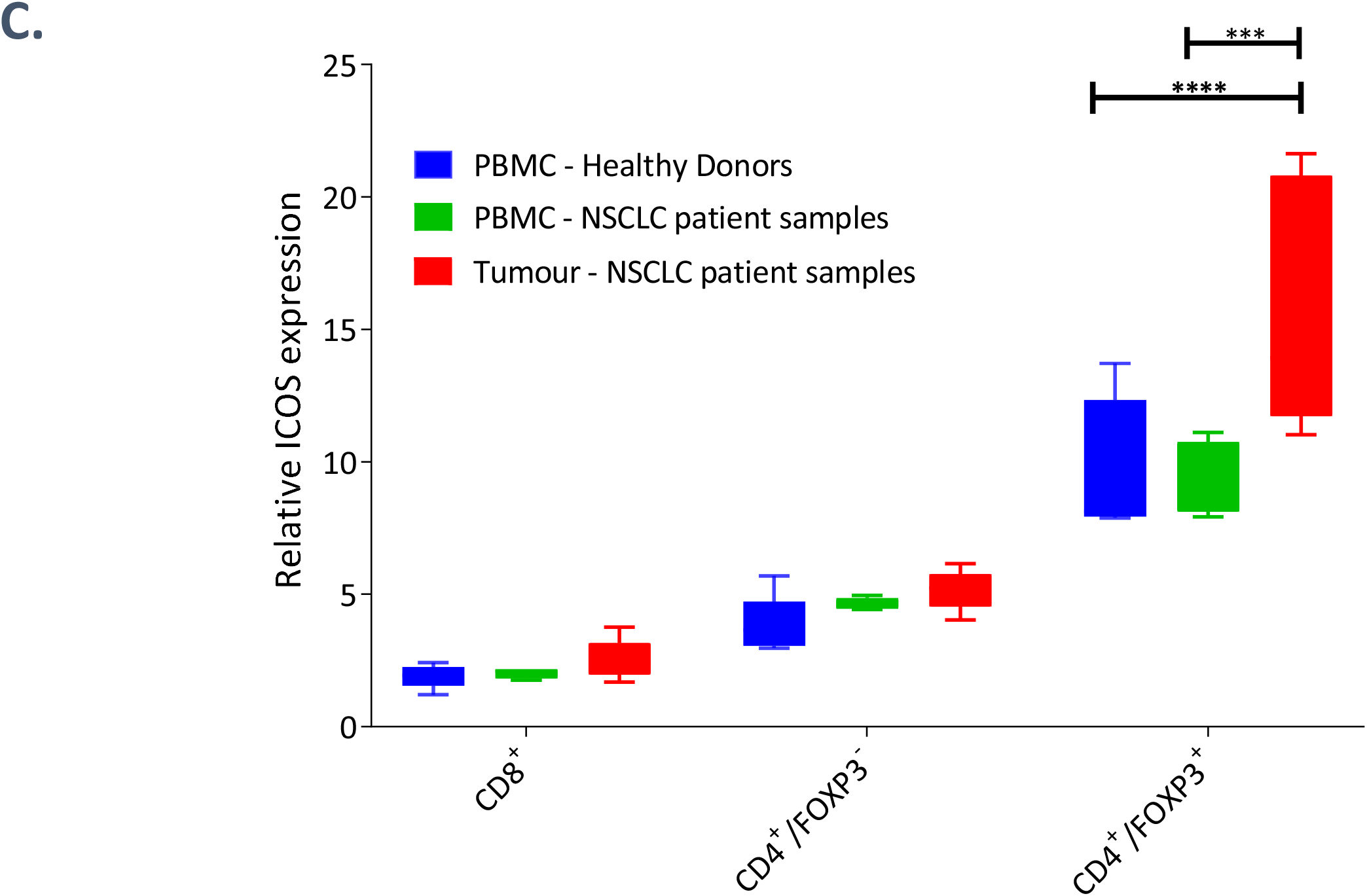
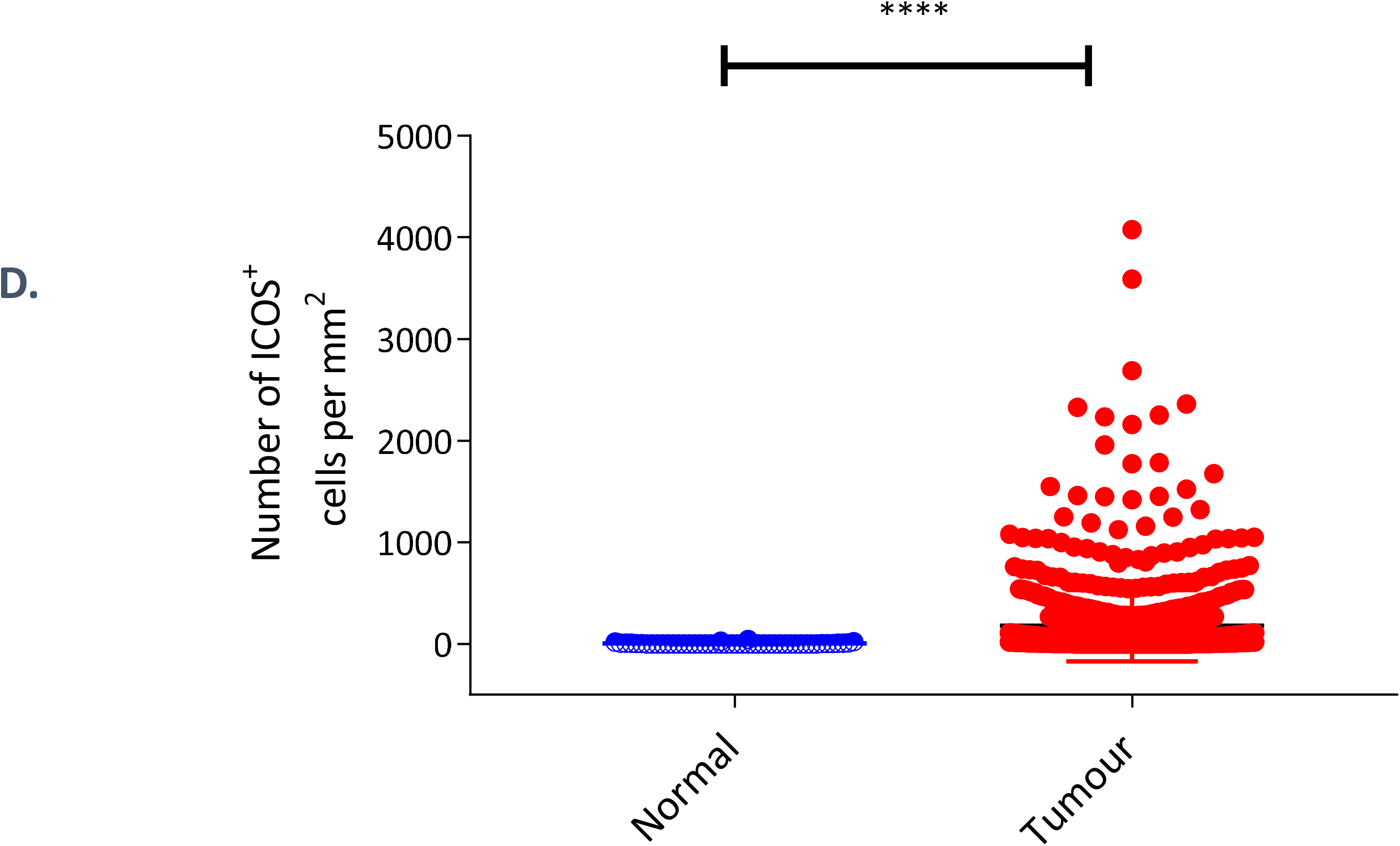
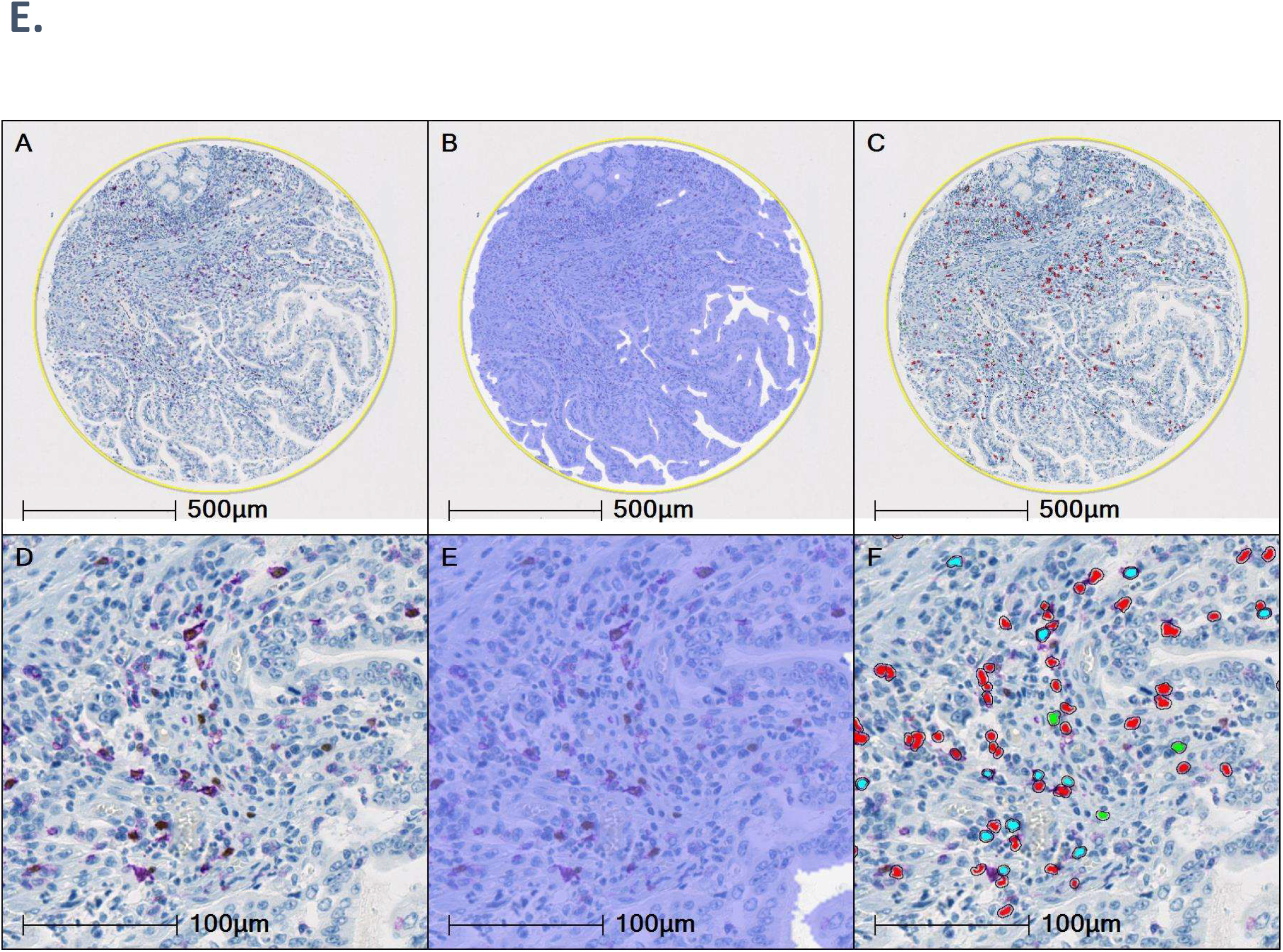
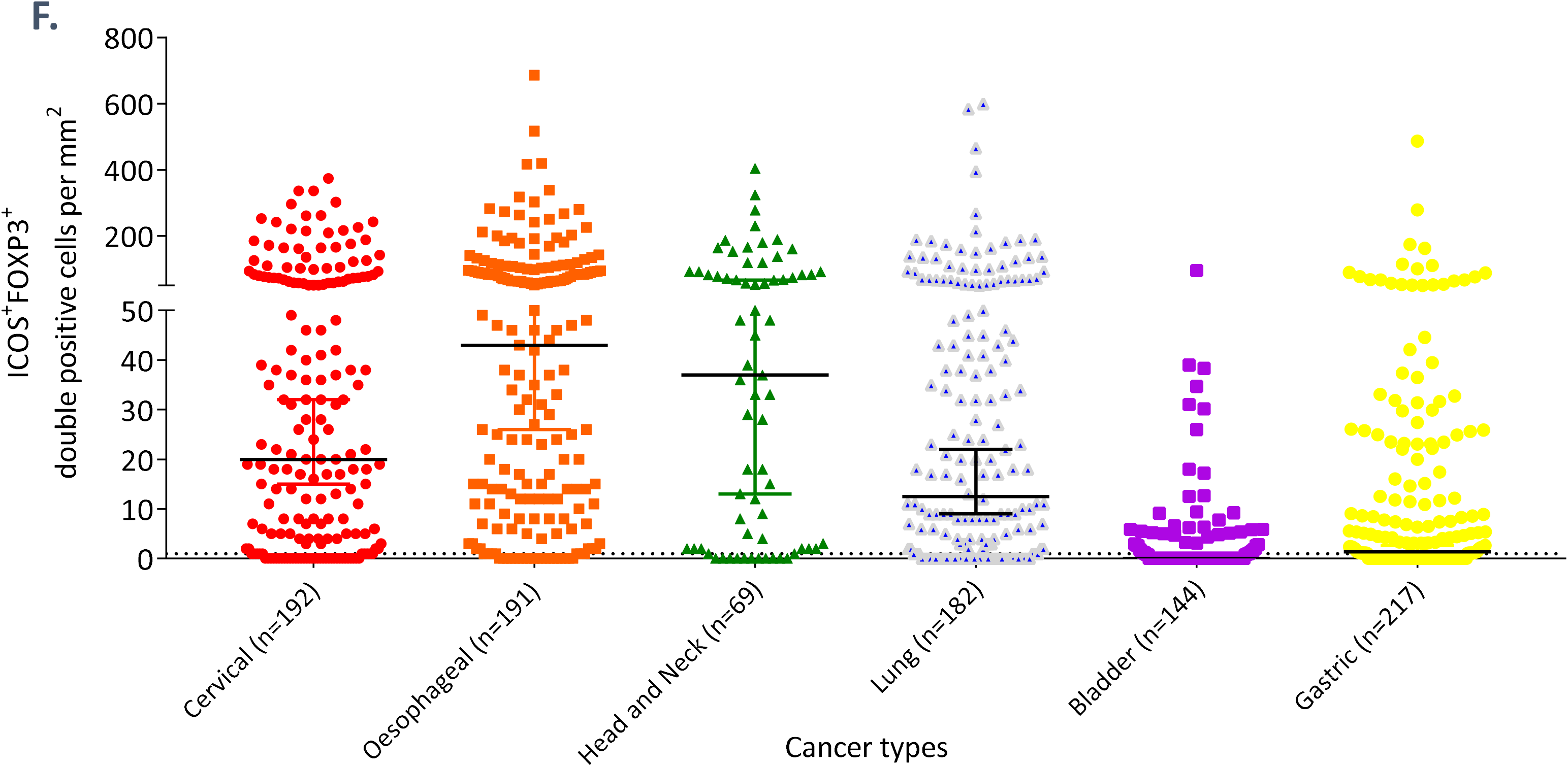
ICOS is highly expressed in cancer T_Regs_. (A) Density plots showing the combined *Icos* and *Foxp3* expression in different solid tumour types from the TCGA datasets. Combined expression was scored using single-sample gene set enrichment analysis (ssGSEA) and density distributions plotted by indications. Stacked histograms are arranged by order of descending median ssGSEA score (marked by vertical ticks). The TCGA standard cancer type abbreviations were used (https://gdc.cancer.gov/resources-tcga-users/tcga-code-tables/tcga-study-abbreviations). (B) Immune response genes were sequenced in each cell. Sequenced cells were then classified into one of 27 immunological subtypes based on their gene signature (cell types and the corresponding gene signatures are described in Supplementary table 1). Heatmap (top) of scaled gene expression in T cells from PBMC (left panel) and tumours (right panel) from NSCLC patient samples (n=5). Genes relevant to T cell lineage and function are presented. Each gene was scaled individually across all cells of the dataset and the mean scaled expression for each of 17 T cell subtypes is presented. Scatter plots (bottom) showing ICOS mRNA expression in T_Reg_, CD4_non-TReg_, CD8 and all other T cells. Each dot corresponds to a single cell. Full lines indicate the mean ICOS gene expression of the total cell compartment and the dashed lines indicate the mode (density peak) of the ICOS+ve cell population. (C) Relative ICOS expression (as determined by flow cytometry analysis) on CD4+, CD8+ and T_Reg_ (CD4+/FOXP3+) in healthy donor PBMCs (n=5), NSCLC tumour samples (n=5) and matched NSCLC patient PBMCs (n=4). *** = P value ≤0.001 and **** = P value ≤0.0001 (2-way ANOVA with Tukey’s multiple comparison). (D) Graph showing the number of ICOS+ve cells per mm^2^ in tumour cores (n= 995) from 6 indications and cores from matching healthy tissues (n=48). ****p≤0.0001 (Mann-Whitney unpaired T test). (E) Example of ICOS/FOXP3 staining of gastric tumour core showing (a); original whole core image with ICOS staining in purple and FOXP3 in brown (b); classifier analysis overlays showing tissue ROI (lilac) and white space (white); (c) cellular analysis overlay showing single FOXP3 positive (green overlay), single ICOS positive (red overlay) and ICOS/FOXP3 dual positive (cyan overlay); (d - f) x20 magnified detailed area (F) Quantification of ICOS/FOXP3 double positive cells per mm^2^ in TMAs from six indications (cervical, oesophageal, head and neck, lung, bladder and gastric) by image digital pathology.

The T cells from these samples (tumours and PBMC) were further analysed by flow cytometry to confirm the levels of ICOS protein on the cell membrane. As shown in Fig. 1C, ICOS expression was higher on the surface of T_Reg_ (CD4^+^/FOXP3^+^) than on the other T cell subsets analysed (CD8^+^ and CD4^+^/FOXP3^-^). Similar to the mRNA levels, ICOS protein levels were higher in the TME than on circulating T cells. This difference between the two compartments was strongly significant (P<0.001) for the T_Reg_ subset. No differences in ICOS expression between PBMCs from healthy donors and NSCLC patients were observed on the T cell subsets analysed. As seen with the human samples, T cell immunophenotyping by flow cytometry analysis revealed that ICOS was also more highly expressed in the TME than in the spleen of CT26.WT syngeneic tumour bearing mice. This analysis also confirmed higher ICOS protein expression levels on T_Reg_ than on CD4/CD8 T_Eff_ cells (Supplementary Fig. 1).

Finally, we assessed tumour tissue microarrays containing cores from oesophageal, head and neck, gastric, lung, bladder and cervical cancer biopsies and carried out immunostaining to detect ICOS. We observed a significant up-regulation of ICOS in the TME as compared with matched healthy tissues (Fig. 1D). In order to confirm the ICOS expression on T_Reg_, tumour cores were co-stained for both ICOS and FOXP3, and the number of cells co-expressing both markers was quantified by digital pathology (Fig. 1E). From this analysis, we further demonstrated the presence of a high number of ICOS^+^ T_Reg_ cells in the TME of the six selected indications, with only bladder cancer demonstrating a lower number of ICOS/FOXP3 double positive cells per mm^2^ (Fig 1F).

Altogether, our data confirmed that ICOS is not homogenously expressed on the different T cell subsets and is strongly induced in the TME, especially on the surface of T_Reg_. In addition, we identified indications with high levels of intratumoural ICOS^+^ T_Reg_ cells and these include oesophageal, head and neck, gastric, lung, and cervical cancers.

### KY1044: a fully-human anti-ICOS IgG1 antibody with both depleting and agonistic functions

The preferentially high expression level of ICOS on intratumoural T_Reg_ cells suggests that the targeting of these cells with an effector enabled anti-ICOS antibody could lead to their preferential depletion (via antibody-dependent cellular cytotoxicity; ADCC) and hence remove their inhibitory effects on the anti-tumour efficacy of ICIs. The level of target expression and the amount of antibody bound to a target are key parameters influencing cell killing by ADCC (*27*). Therefore, it is expected that ICOS^Low^ cells (e.g. CD8^+^ T_Eff_ cells) will be less sensitive to depletion than ICOS^High^ T_Reg_ but yet could be sensitive to co-stimulatory potential via ICOS signalling if the antibody also harbours agonistic properties (via clustering through Fc*γ*R receptors) (*28, 29*). With this in mind, we identified an antibody called KY1044. KY1044 is a human monoclonal IgG1 (effector enabled) antibody that was generated in atransgenic mouse line in which the endogenous *Icos* gene was knocked-out. Lack of endogenous ICOS expression was aimed to facilitate the generation of a mouse cross reactive antibodies. KY1044 binds ICOS from different species (human, Cynomolgus monkey, rat and mouse) with strong affinity (a KD below 3 nM for the Fab).

In vitro, KY1044 ADCC potential was tested in a series of cell-based assays using established cell lines as well as primary T cells and natural killer (NK) cells. The ability of KY1044 to induce ADCC was first assessed using a luciferase reporter assay. This assay quantifies the activation of FcγRIIIa signalling in response to KY1044 bound onto ICOS-expressing CHO cells. As shown in Fig. 2A, binding of KY1044 to human ICOS-expressing CHO cells significantly induced the luciferase signal with an average EC_50_ of 0.15 nM (n=3). A similar EC_50_ was obtained when using CHO cells expressing mouse ICOS (0.53 nM, n=3), rat ICOS (0.48 nM, n=3) or cynomolgus monkey ICOS (0.22 nM, n=3). No luciferase signal was induced with the human IgG1 isotype control. To validate further the ADCC potential of KY1044, we used purified human NK cells as effector cells. When incubated with KY1044 and CEM cells expressing human ICOS (5:1 Effector:Target cell ratio), these NK cells induced potent ADCC-mediated killing with an average EC_50_ of 5.6 pM (n=8, Fig. 2B). Altogether, the data confirmed that KY1044 has the ability to trigger ADCC (and probably ADCP) of ICOS^High^ expressing cells though engagement with FcγRIIIa.

**Figure 2:**
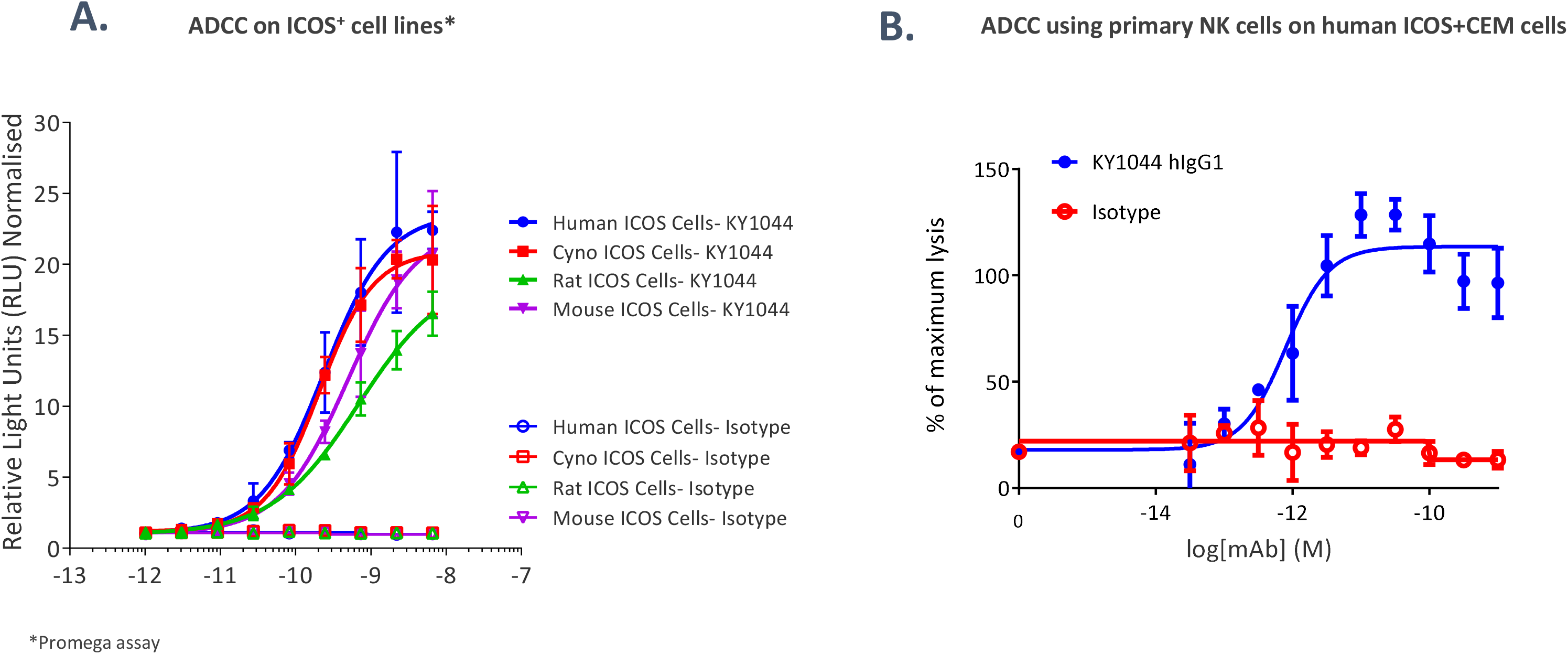
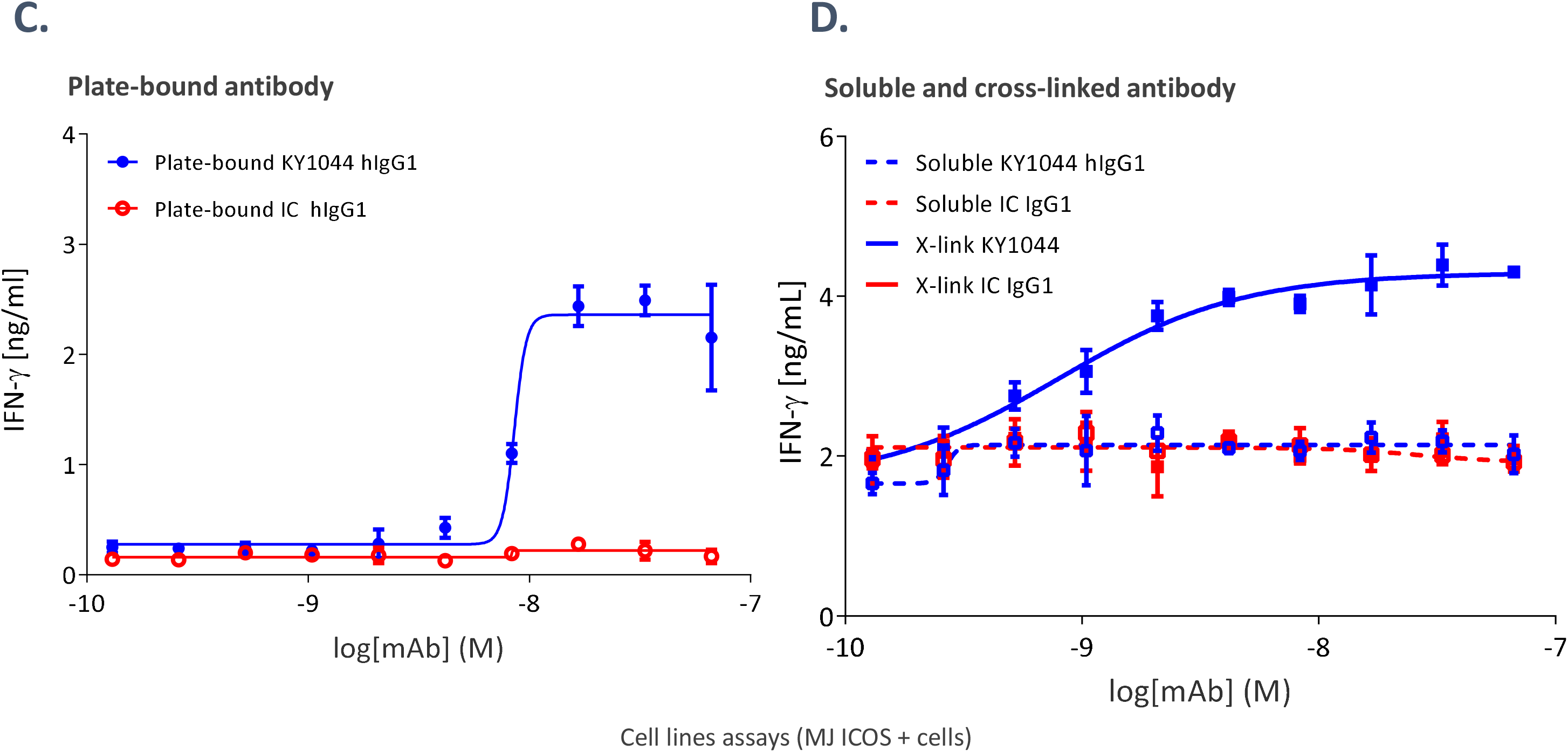
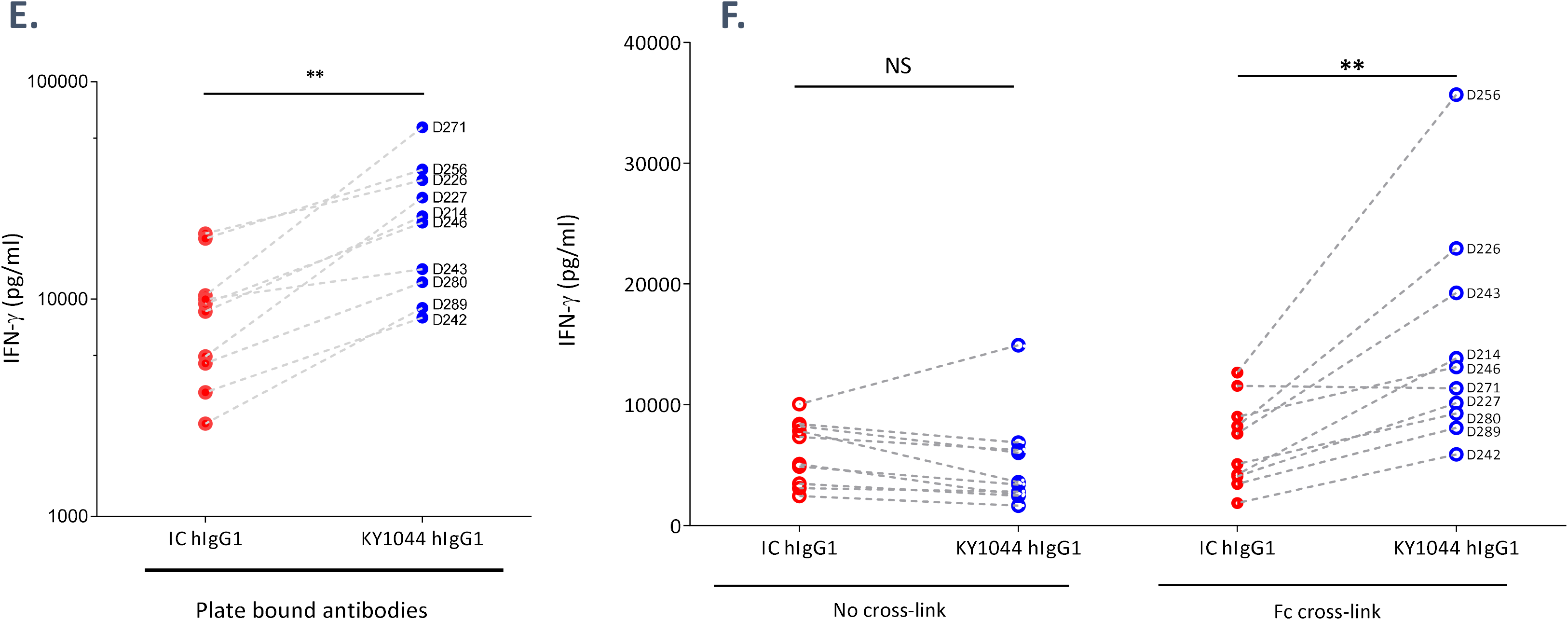
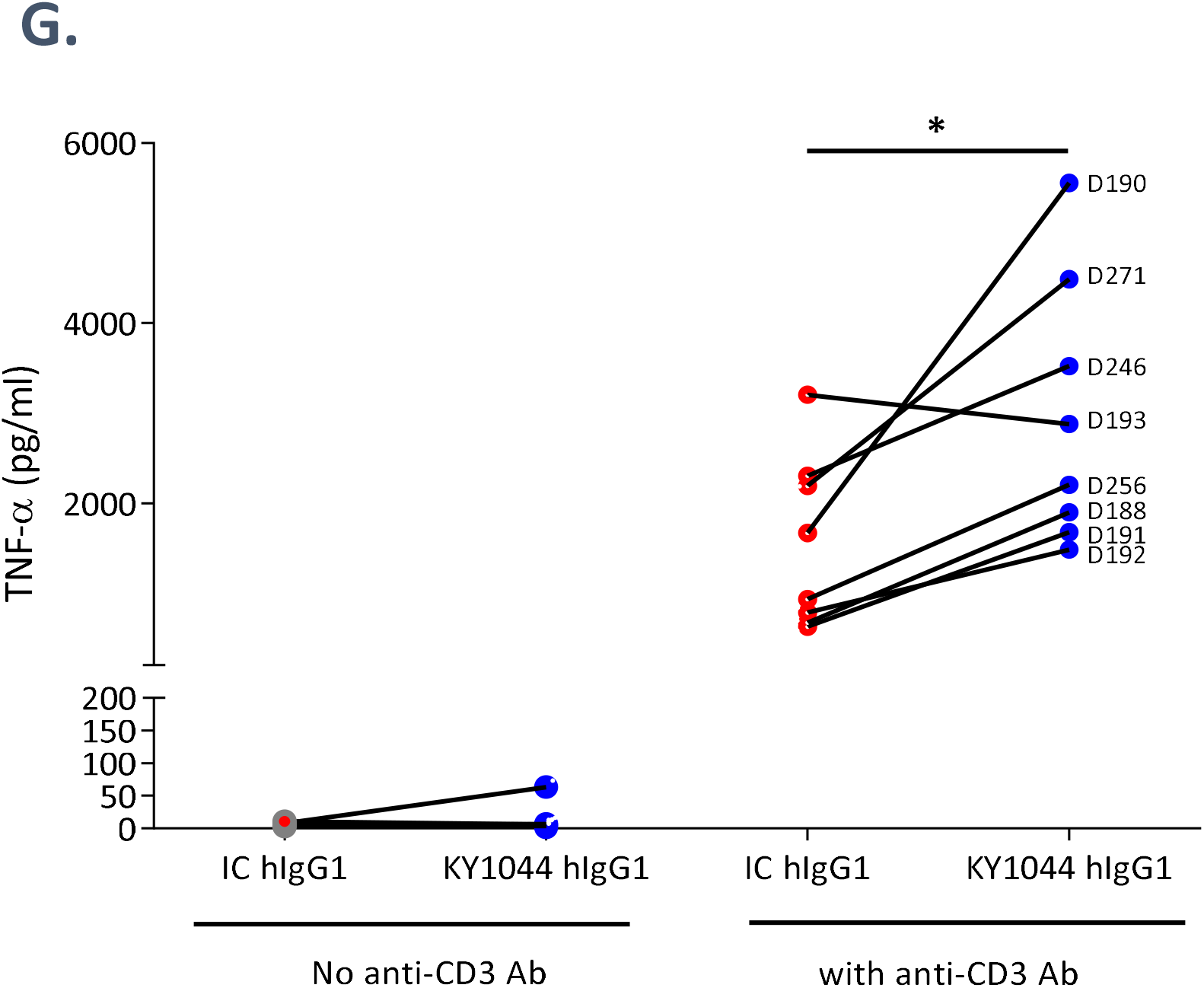
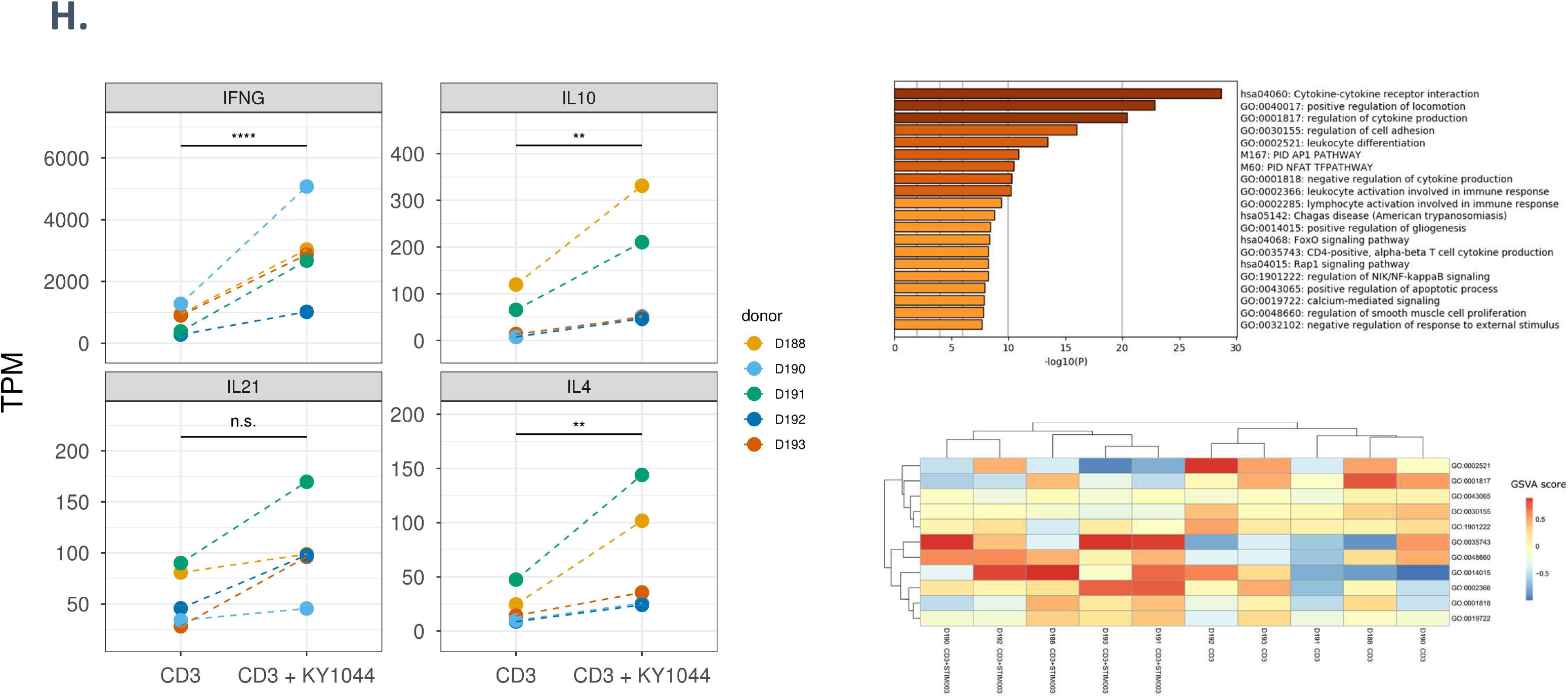
KY1044 triggers ADCC and has co-stimulatory agonistic properties in ICOS positive T cells (A) KY1044 induces FcgRIIIA signalling in “effector” cells when cocultured with ICOS positive cells. Serial dilutions of KY1044 hIgG1 and the isotype control were incubated O/N with engineered Jurkat effector cells containing the reporter NFAT-luciferase and FcγRIIIa and CHO cells expressing ICOS from different species. (B) KY1044 induces ADCC in a human primary NK cells assay. Serial dilutions of KY1044 or isotype control were co-incubated with ICOS CEM target cells and NK for 4 hours. ADCC activity was quantified by measuring Specific Dye Release (mean of triplicates±SEM). (C) and (D) Plate bounds and cross-linked soluble KY1044 induces the release of IFN-γ from MJ cells. The concentration of IFN-γ was assessed in the supernatant of MJ cultured for 72hrs in the presence of serial dilutions of plate bound and soluble antibodies with or without the addition of an Fc cross-linking anti-human F(ab’)₂ Fragments (D). (E) and (F) Levels of IFN-γ induced in human primary T-cell activated with anti-CD3/CD28 (to induce ICOS) and cultured with either 5 μg/mL of plate-bound KY1044 or the isotype control (IC) IgG1 (E, n=10) or with either 15μg/mL of soluble KY1044 or soluble IC, with or without the addition of an Fc cross-linking anti-human F(ab’)₂ Fragments (F, n=10). (G) Cytokine production by KY1044 requires anti-CD3. Levels of TNF-α in T-cell culture with 5 μg/mL of KY1044 or IC IgG1 (plate-bound, n=8) in the presence or not of anti-CD3. * p<0.05, ** p<0.01 (Wilcoxon statistic test). (H) RNA sequencing analysis comparing T cell stimulation with either KY1044 + anti-CD3 or control (anti-CD3 only) antibodies. Genes reported to be downstream of ICOS signalling show a pattern of upregulation following KY1044 co-stimulation, with significant p-values obtained for IFNG, IL10 and IL4 following 6 hrs of stimulation (left). Metascape pathway enrichment analysis of the differentially expressed genes from the 6-hr time point. Enriched gene sets are ranked based on the significance of their p-values (top right). Heatmap showing unsupervised clustering of samples based on the expression levels of the top GO terms from the Metascape enrichment analysis. KY1044 stimulated samples show specific gene ontology modules that segregate them from control stimulated samples (bottom right).

The agonistic potential of KY1044 was assessed using the MJ [G11] CD4^+^ cell line, which endogenously expresses ICOS and does not require addition of a primary stimulatory signal (e.g. TCR engagement) for cytokine production. KY1044 was either pre-coated on culture plates (mimicking cross-presentation/clustering) or used in solution with or without addition of a secondary cross-linking antibody to cluster KY1044. KY1044 did not increase proliferation of ICOS^+^ T cells. However, we demonstrated agonism by quantifying pro-inflammatory cytokines such as IFN-γ and TNF-α released in the media. While soluble KY1044 did not induce IFN-γ production, plate-bound and cross-linked KY1044 effectively induced IFN-γ secretion in a concentration-dependent manner with an EC_50_ of 10.5 nM (±2.7 nM; n=2) and 0.50 nM (±0.18 nM, n=3), respectively (Fig. 2C and 2D). The agonism potential of KY1044 was also confirmed using purified primary T cells. Unlike MJ cells, T cells from PBMC were pre-activated with anti-CD3/CD28 beads (to induce ICOS expression; Supplementary Fig. 2) and cultured in the presence of KY1044 coated onto plates or presented in solution. As observed with the MJ cell line, plate-bound (Fig. 2E) and cross-linked (Fig. 2F) KY1044 induced IFN-γ production in primary cells whilst soluble KY1044 did not. The levels of IFN-γ released in response to KY1044 agonism were variable between donors. Despite this variability, KY1044-dependent IFN-γ secretion was significantly higher than for the isotype treated cells in both plate-bound (2.4 ±0.5-fold at 5 μg/ml mAb, p<0.01) and cross-linked assays (2.5±0.2-fold at 15 μg/ml mAb, p<0.01). Similar data were obtained with induction of TNF-α(Fig. 2G). Importantly, a three-step stimulation-resting-costimulation experiment confirmed that TCR engagement was essential for KY1044 agonism. Indeed, the up-regulation of cytokines (IFN-γ and TNF-α) by ICOS^+^ T cells did not occur without concomitant TCR activation (Fig. 2G).

Finally, the direct agonistic potential of KY1044 to costimulate a pool of purified CD3^+^ T cells was confirmed using whole transcriptome RNA sequencing following 6-hours of combined anti-CD3 (to induce ICOS) and KY1044 stimulation. In line with previous reports (*14*) and our own cytokine measurements, we observed upregulation of key cytokine genes, most notably *Ifng, IL10 and IL4* (Fig. 2H), Gene set enrichment analysis of the transcriptome confirmed that KY1044-dependent ICOS agonism induces genes involved in cytokine-cytokine receptor interactions as well as both leukocyte and lymphocyte activation, which did not seem to be significantly dependent on cell proliferation (Fig 2H). In fact, no significant change in genes associated with proliferation and cell cycle progression was observed. The analysis also showed an enrichment of genes involved in cell locomotion, adhesion and differentiation(*30*). This finding reflected our observations of MJ [G11] CD4^+^ morphology changes following long term anti-ICOS (C398.4A or KY1044) plate bound stimulation (Supp. Fig 2C and D). Even though ICOS signalling was previously reported to depend on the PI3K/AKT/mTOR axis (*31*), for KY1044 agonism, we mainly noticed an overrepresentation of genes downstream of the nuclear factor of activated T cells (NFAT) which is in agreement with recent findings (*32*). NFAT-dependent ICOS agonism was confirmed using another anti-ICOS (C398.4A) in a series of NFAT luciferase reporter cell lines also expressing ICOS with or without CD3ζ (Supp Fig 2E).

Altogether the data presented here demonstrated that KY1044 has a dual mechanism of action with both depleting and agonistic co-stimulatory properties.

### KY1044 (mIgG2a) monotherapy blocks tumour growth in lymphoma/myeloma syngeneic tumour models

KY1044 cross-reactivity to mouse ICOS facilitated the pre-clinical *in vivo* mouse pharmacology work for which the antibody was reformatted as a mouse IgG2a (the effector enabled format in mouse (*33*)). Outside of the solid tumour TCGA datasets (Fig. 1), we observed that ICOS expression was high in Diffuse Large B-Cell Lymphoma (DLBCL, Supplementary Fig. 3). Based on the known role of ICOS/ICOS-LG signalling in the generation and maintenance of germinal centres (*10, 34*) and on previously reported anti-tumour efficacy by anti-ICOS antibodies in lymphoma (*35*), therefore we assessed the anti-tumour efficacy of KY1044 mIgG2a in the ICOS-LG^+^ A20 tumour lymphoblast B cell model (*36*). Mice were dosed (2qw for 3 weeks at 10 mg/kg) starting from 6 days post tumour cell implantation with saline or with KY1044 mIgG2a. Both treatments were well tolerated, and no bodyweight loss was observed. Importantly, when compared to the control group, KY1044 mIgG2a resulted in an effective anti-tumour efficacy, with more than 90% of animals being free from measurable tumours at the end of the study (day 42, Fig. 3A). In addition, efficacy was confirmed (albeit to a lesser extent) in another B cell-derived syngeneic model, the J558 plasmacytoma model. As observed in the A20 model, KY1044 exhibited a clear anti-tumour efficacy, with around 70% (5 out of 7, Fig. 3B) of the KY1044 mIgG2a-treated mice being tumour free at the end of the study. Monotherapy response in non B-cell-derived tumour syngeneic models was also investigated. These models included models of haematological malignancies (T-cell lymphoma EL4) and models of solid tumours (the colorectal cell lines CT26.WT and MC-38 and the melanoma cell line B16.F10). Overall, the monotherapy response was absent or low in all these models. Mice harbouring CT26.WT or MC-38 tumours showed limited monotherapy anti-tumour response with only some tumour growth delay (supplementary Fig. 4). Altogether, our preclinical in vivo mouse tumour studies demonstrated that KY1044 mIgG2a displayed mild anti-tumour response in CT26.WT and MC-38 mouse colon cancer model but was highly effective as a monotherapy in B cell-derived mouse tumour models.

**Figure 3:**
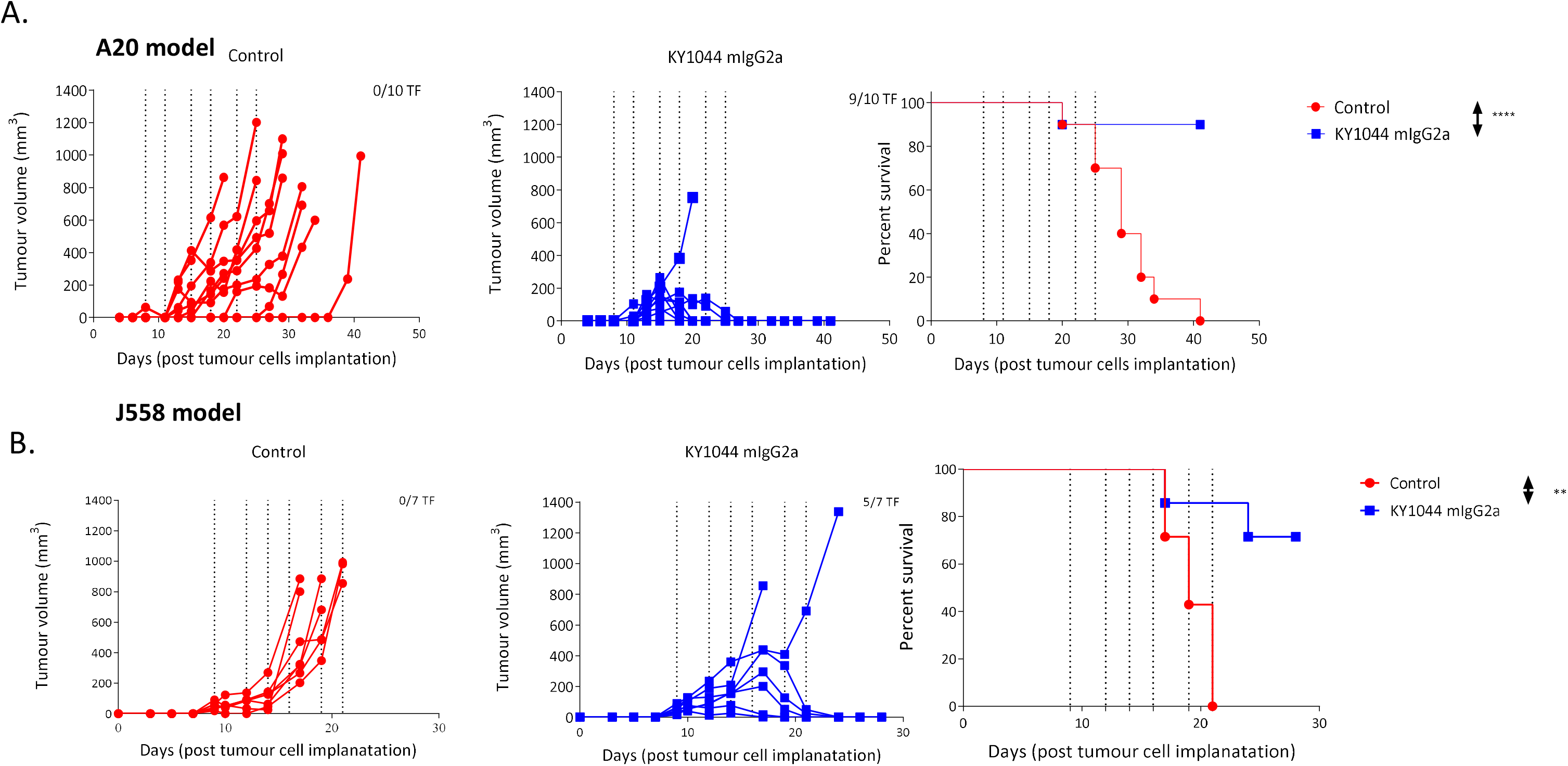
KY1044 mIgG2 monotherapy triggers anti-tumour efficacy in the A20 and J558 syngeneic tumour models (A) Spider plots showing individual mouse tumour volumes from BALB/c mice (n=10 per group) harbouring subcutaneous A20 tumours. Tumour bearing mice were dosed i.p with 200 µg of KY1044 mIgG2a or 200 µl saline starting from day 8 post tumour cell implantation. (B) Spider plots showing individual mouse tumour volumes from BALB/c mice (n=7 per group) harbouring subcutaneous J558 tumours. Tumour bearing mice were dosed i.p with 60 µg of KY1044 mIgG2a or 200 µL saline starting from day 9 post tumour cell implantation. For both experiments the dosing days are indicated by vertical dotted lines. The Survival curves depicting the control and KY1044 mIgG2a groups are shown on the right end side. ** P<0.01, **** P<0.0001 (Statistics calculated using Log-rank (Mantel-Cox) test. Data are representative of at least 2 experiments.

### KY1044 (mIgG2a) synergises with anti-PD-L1 in models resistant to both monotherapies

As discussed above, only around 10% of the treated mice harbouring CT26.WT tumours responded to KY1044 mIgG2a monotherapy. Likewise, CT26.WT tumours are known to poorly respond to anti-PD-L1 monotherapy (*37*). Since anti-ICOS and the anti-PD-L1 antibodies act on different but complementary immune pathways, we assessed whether the combination could improve the anti-tumour response. These studies (Fig. 4A) demonstrated a complete anti-tumour response in the majority (70 %) of mice treated with the KY1044 mIgG2a (equivalent of 3 mg/kg) and anti-PD-L1 (equivalent of 10 mg/kg) combination. In other in-house experiments, this response rate ranged from 50 to 90% of treated mice. Of relevance, when KY1044 was reformatted as a mouse IgG1 (low depleting potential in mouse) and combined at the same dose with anti-PD-L1, the resulting anti-tumour efficacy was weaker than the one observed with the KY1044 mIgG2/anti-PD-L1 combination, thus arguing for the contribution of ICOS-mediated depletion to the stronger anti-tumour efficacy seen in the CT26.WT model (Supplementary Fig 6A).

**Figure 4:**
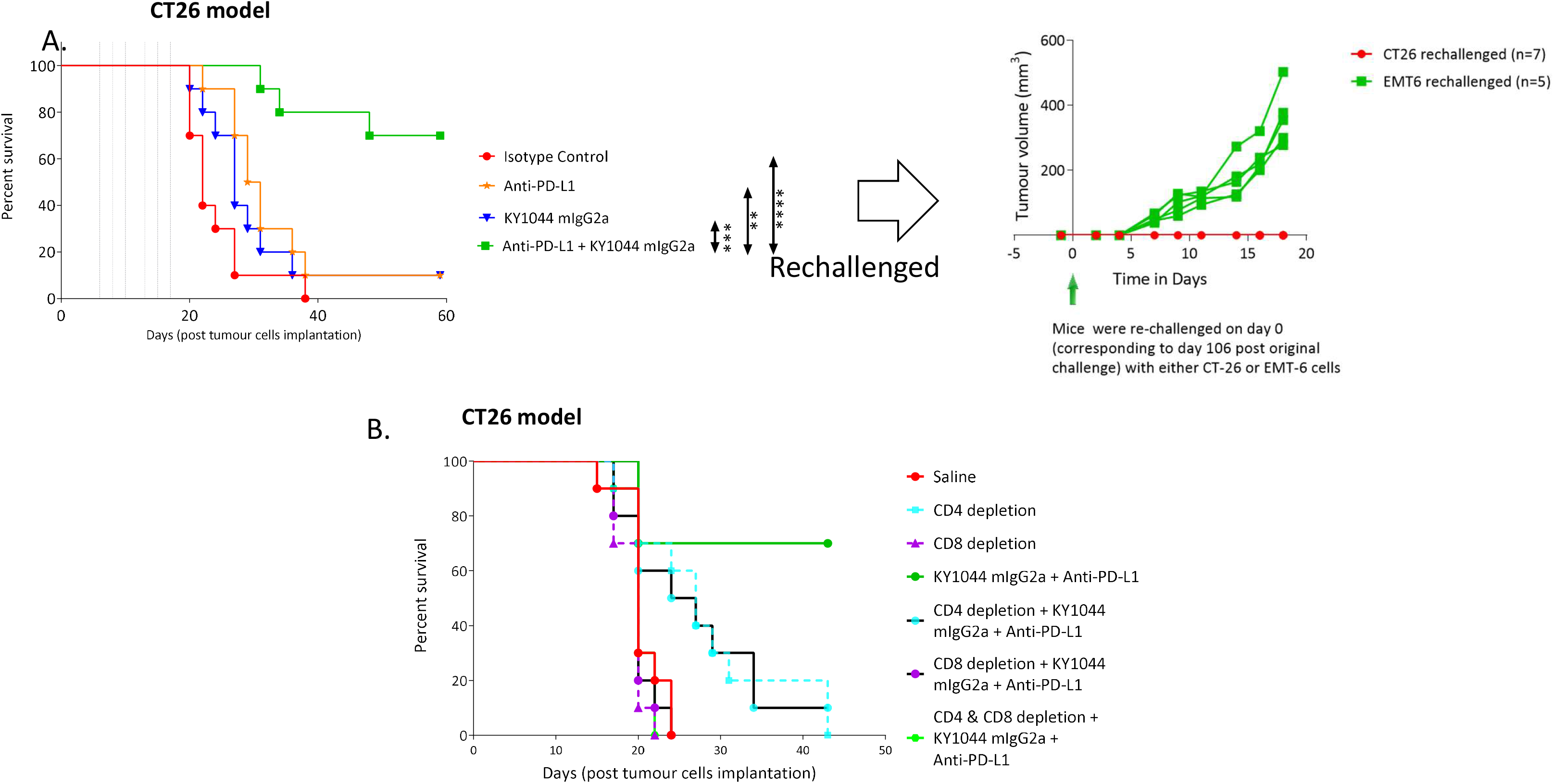
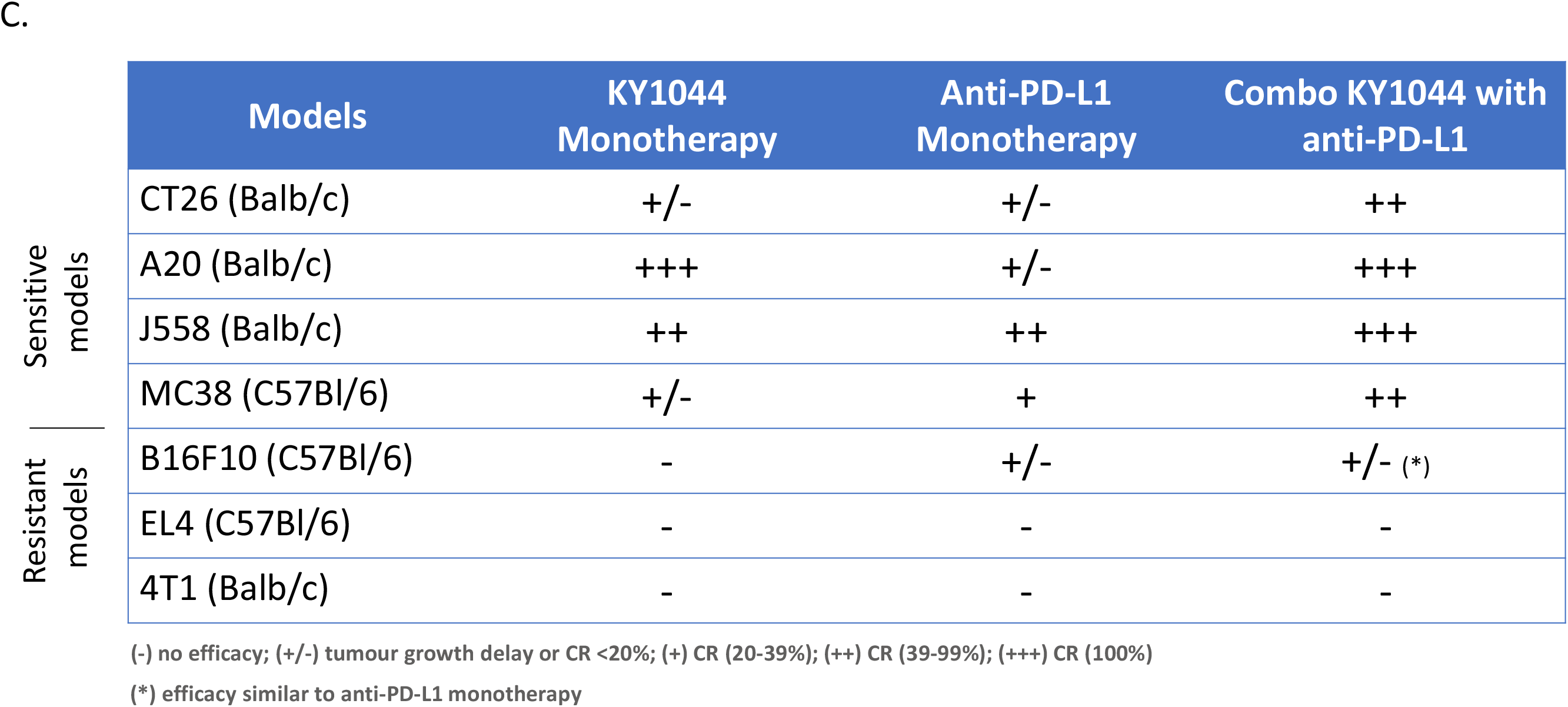
KY1044 mIgG2a synergises with anti-PD-L1 to promote durable anti-tumour immune response in model of solid tumours. (A) Survival curves of mice harbouring CT26.WT tumours and treated with control, KY1044 mIgG2a monotherapy at 60 µg/dose, anti-PD-L1 monotherapy 200 µg/dose and the combination of KY1044 mIgG2a and anti-PD-L1 at 60 and 200 µg/dose, respectively (Left panel). The dosing days are indicated by vertical dotted lines. ** P<0.01, *** P<0.001 and **** P<0.0001 (Statistics calculated using Log-rank Mantel-Cox test). The data is representative of 6 independent experiments. Tumour-bearing mice cured of CT26.WT tumours were randomly allocated to two groups and rechallenged with either CT26.WT or EMT6 tumour cells (Right panel). Tumour growth was only observed for EMT6 tumours but not CT26.WT tumours. (B) The anti-tumour response to the combination is primarily dependent on the presence CD8 (and to a lesser extent on CD4) T Cells. Mice implanted with CT26.WT cell (n=10 per group) were depleted of CD8 and/or CD4 T cells and treated with saline or with KY1044 mIgG2a and anti-PD-L1 combination as described in panel (A). Data is representative of two independent experiments. (C) Table summarising the antitumour efficacy of KY1044 mIgG2a monotherapy, anti-PD-L1 monotherapy and the combination in different syngeneic models.

Notably, the mice that survived the first CT26.WT challenge in response to the KY1044 mIgG2a/anti-PD-L1 treatment were shown to be resistant to a CT26.WT re-challenge but sensitive to the growth of the EMT-6 tumour cell line (Fig. 4A). This response confirmed that the combination therapy not only induced an anti-tumour response but also generated a tumour antigen specific memory response in these “cured” mice. Finally, we demonstrated that the response to the KY1044 mIgG2a and anti-PD-L1 combination was primarily dependent on CD8^+^ T cells, with the improved survival fully abrogated when CD8^+^ T cells were depleted (Fig. 4B and supplementary Fig. 5). Conversely, addition of an anti-CD4 depleting antibody to the combination only reduced but did not fully prevent the anti-tumour response. Finally, pre-treatment with anti-CD4/CD8 prevented the response to the combination to a similar level to that observed with the anti-CD8 depleting antibody alone (Fig. 4B). Altogether, the data demonstrated that the co-administration of KY1044 mIgG2a and anti-PD-L1 mAbs triggers a strong and durable anti-tumour immune response in the CT26.WT syngeneic tumour model. Interestingly, the combination of anti-PD1 antibody (clone RMP1-14) with KY1044 was poorly effective in this model (supplementary Fig. 6B).

The KY1044 mIgG2a and anti-PD-L1 combination was also tested in additional BALB/c and C57Bl/6 syngeneic tumour models (Fig. 4C). Although the J558 model already responded well to either the anti-ICOS or the anti-PD-L1 monotherapy, a 100% response was achieved when combining both agents in this model. Similarly, the combination produced an effective response in the C57Bl/6 MC-38 model (with 62.5% complete response in comparison with 0% complete response observed with the respective monotherapies; Supplementary Fig. 7). It should be noted that not all syngeneic tumours responded to the KY1044/anti-PD-L1 combination. For example, no response was observed in the B16F10, 4T1 and EL4 tumour models (Fig. 4C). Collectively, our preclinical tumour efficacy studies clearly demonstrate that the co-targeting of ICOS and PD-L1 results in a strong synergistic effect in selected tumour models, including those in which both monotherapies are poorly effective.

### KY1044 depletes ICOS^High^ cells *in vivo* in mice and non-human primates

In order to assess the mechanism of action of KY1044 *in vivo,* we first conducted pharmacodynamic (PD) studies in the CT26.WT syngeneic tumour model. In this experiment, mice harbouring CT26.WT tumours were dosed twice with a dose response of KY1044 mIgG2a (ranging from 0.3 to 10 mg/kg) on day 13 and 15 post tumour cell implantation (Fig. 5A). The T cell content in the tumours and the spleens were then analysed by flow cytometry 24 hours after the second dosing. Although KY1044 monotherapy was not sufficient to induce strong anti-tumour efficacy in the CT26.WT model (Fig. 4A), the PD study shows a strong effect on the tumour immune contexture at all doses tested. KY1044 mIgG2a was strongly associated with intratumoural T_Reg_ cell depletion, even at doses as low as 0.3 mg/kg. Noteworthy, this decrease in T_Reg_ was significant for doses equal or superior to 1 mg/kg. T_Reg_ depletion was associated with an increase in the CD8^+^ T_EFF_ to T_Reg_ cell ratio (known to be associated with improved response to ICIs) in the TME at all tested doses (Fig. 5B). However, the highest dose of 10 mg/kg showed a lower (yet superior over saline control) CD8^+^ T_EFF_ to T_Reg_ cell ratio than the one resulting from the 1 and 3 mg/kg doses. This decrease in the ratio at the highest dose may have been caused by the depletory effect (albeit not significant) of CD8^+^ T cells (Supplementary Fig. 8A) in response to KY1044 mIgG2a. Noteworthy, when a similar analysis was performed on the spleens of treated mice, no depletion of T_Reg_ and no changes in the CD8^+^ T_EFF_ to T_Reg_ cell ratio were observed (Fig. 5C and Supplementary Fig. 8B) probably due to lower ICOS expression the spleen (Supplementary Fig. 1). We repeated similar PD experiment by dosing once CT26.WT tumour bearing mice with KY1044 mIgG2a. The tumours were then isolated for immunophenotyping up to seven days post treatment (Supplementary Fig. 8C). In this experiment, depletion of T_Regs_ was observed after a single dose of 0.3mg/kg, however at this dose, the incidence of T_Regs_ in the TME was back to the level observed in the control group. A significant and long-lasting T_Regs_ depletion was observed at higher dose (3 and 10mg/kg) and as well as an improvement of the CD8 to T_Regs_ ratio which was significant at intermediate dose. Altogether this data suggests that KY1044 mIgG2a preferentially depletes ICOS^high^ T_Reg_ cells in the TME probably due to the higher ICOS expression described in the tumour (Supplementary Fig.1).

**Figure 5:**
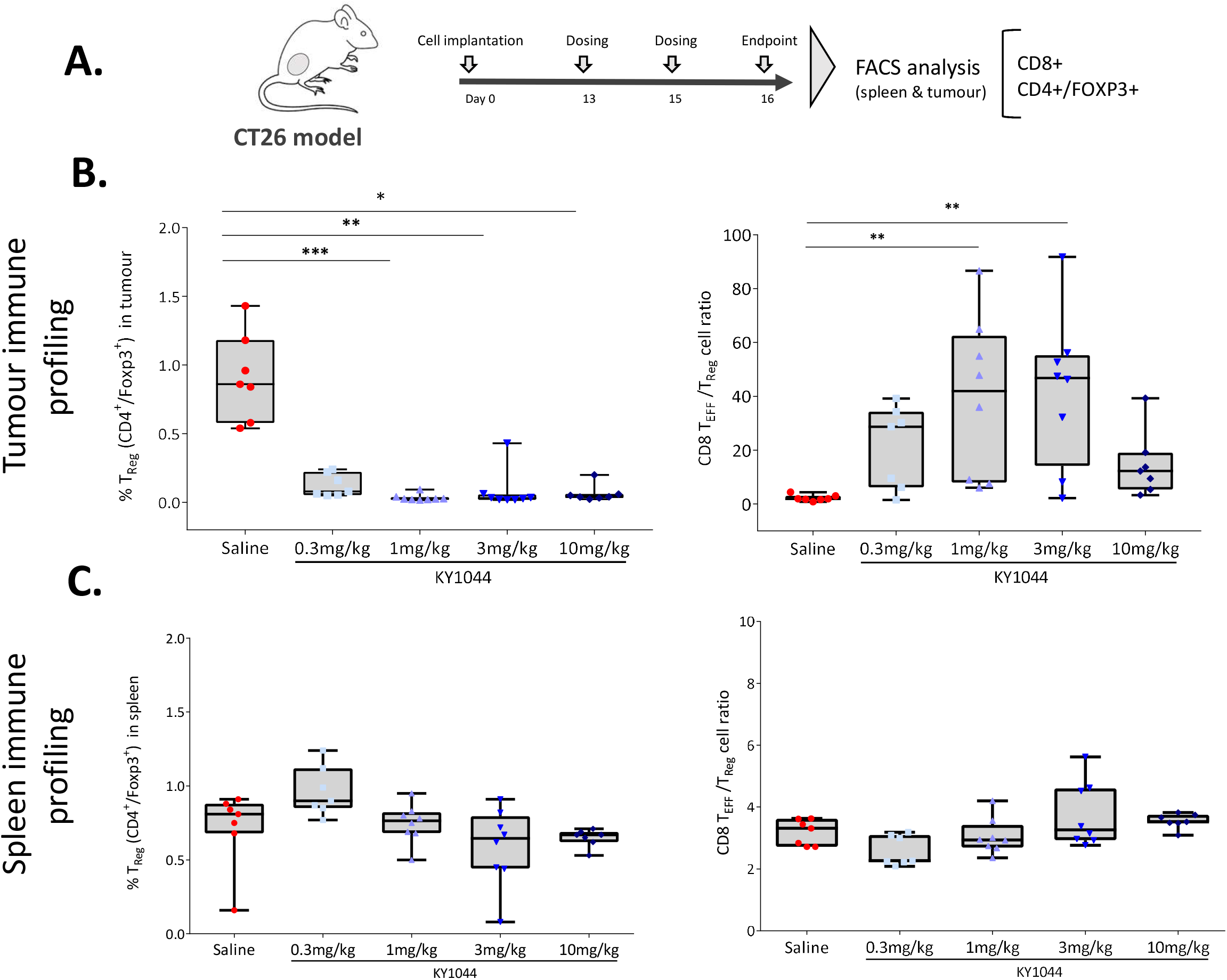
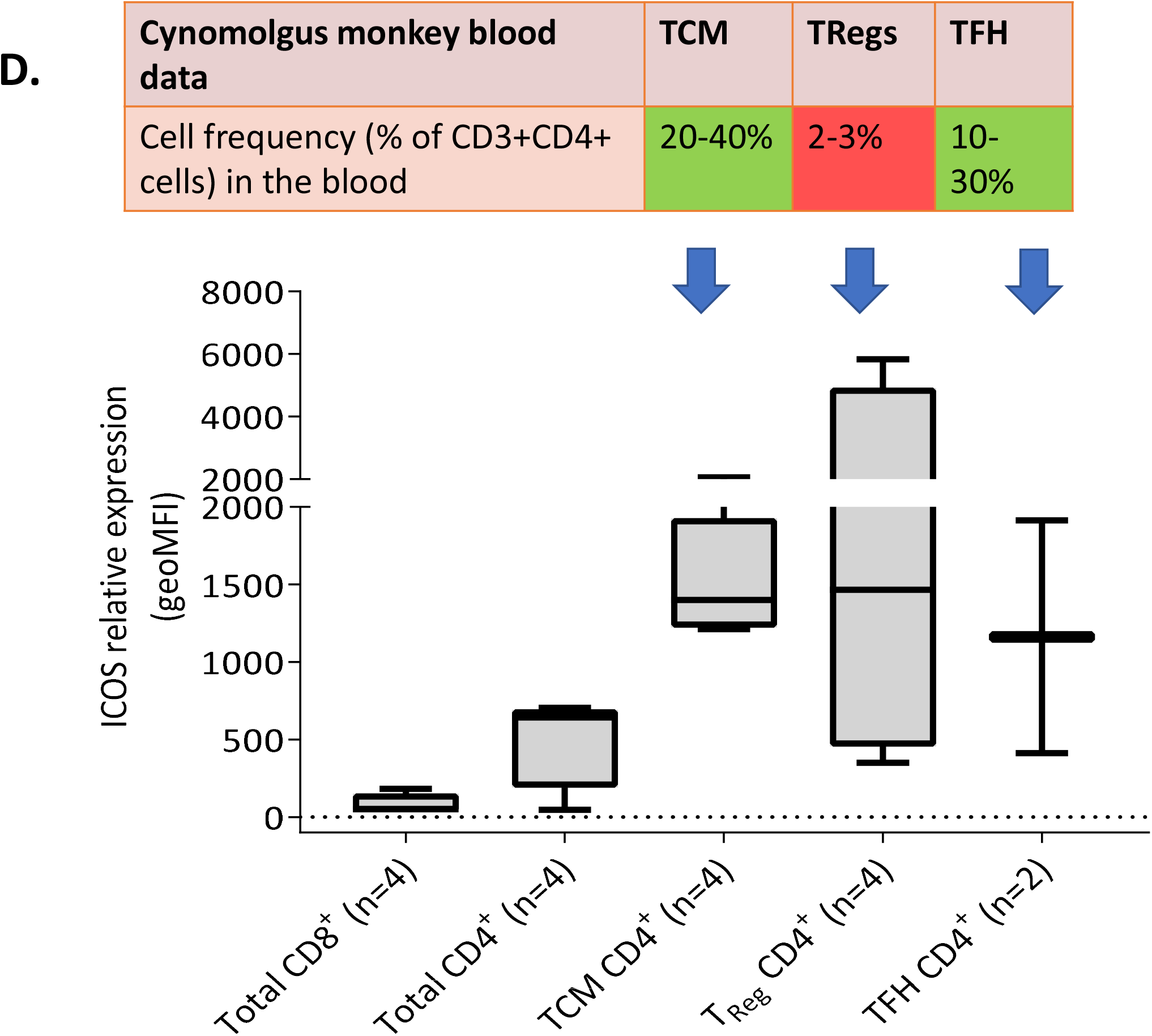
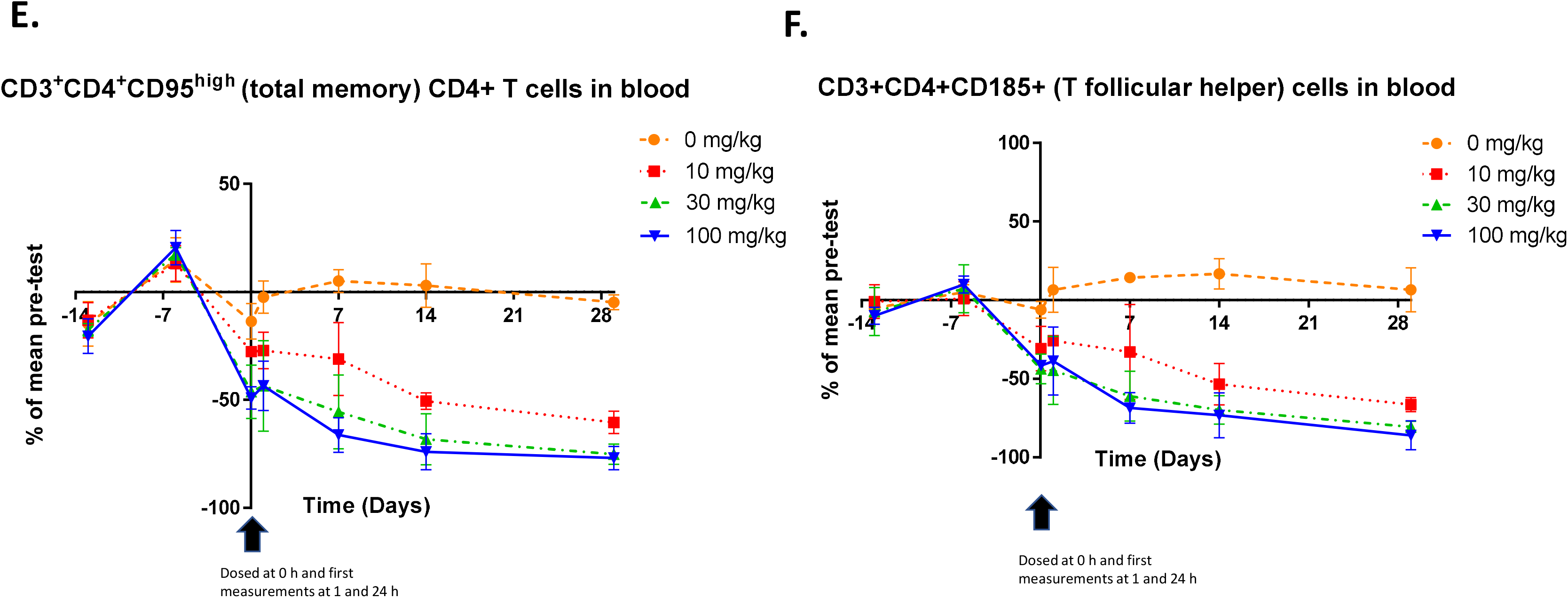
KY1044 mIgG2a preferentially depletes ICOS^high^ T_Reg_ cell *in vivo*. (A) In vivo experimental protocol of the pharmacodynamic study in the CT26.WT tumour bearing BALB/c mice. Mice were dosed twice with saline or different doses of KY1044 mIgG2a (ranging from 0.3 to 10 mg/kg) on days 13 and 15 and the tumour and spleen were harvested on day 16 and the immune cells analysed by flow cytometry. KY1044 depletes T_Reg_ and increase the CD8+ T_EFF_ to T_Reg_ cell ratio in the tumour microenvironment (B) but not in the spleen (C) of treated mice (n=7/8 mice per groups). * p<0.05, ** p<0.01 and ***p<0.001. (D) Relative expression of ICOS and frequency of T cells subsets in the blood of NHP (the number of NHP used for each analysis are indicated in brackets). ICOS is highly expressed on T_Reg_ cell followed by CD4 memory (T_M_) and T follicular helper cells (T_FH_). NHP CD8 T cells express low levels of ICOS. (E) (F) and (G) Graphs showing the changes (vs baseline pre-treatment) of total memory CD4 T cells and follicular T Helper cells in the blood of NHP at different timepoints following a single dose KY1044 hIgG1 at 0, 10, 20 and 100 mg/kg (n=3 NHP per groups)

As for human immune cells (Fig.1), we confirmed that ICOS was expressed in cynomolgus monkeys on CD4^+^ memory T cells (T_M_ defined as CD3^+^/CD4^+^/CD95^high^/CD28^-/Dim/+^), CD4^+^ follicular T helper cells (T_FH_ defined as CD3^+^/CD4^+^/CD185^+^) and on T_Reg_ cells (defined as CD3^+^/CD4^+^/CD25^+^FoxP3^+^) (Fig. 5D). As previously shown for other species, circulating NHP CD8^+^ T cells showed the lowest levels of ICOS expression. We assessed the pharmacodynamic effects of KY1044 in cynomolgus monkeys after repeated weekly i.v. administration at dose levels of 0, 10, 30 and 100 mg/kg for 4 weeks (5 doses). Exposure and full occupancy of ICOS on circulating CD4+ T cells was maintained at all dose levels throughout the study (Supplementary Fig. 9A-B). Detailed immunophenotyping of different T cell subsets was performed by flow cytometry at various times up to 29 days after the first dose. This analysis indicated that there was a marked decrease in T_M_ and T_FH_ cells in peripheral blood (Fig. 5E-F) which was apparent soon after dosing. Absolute T_M_ counts at the end of the study (28 days post dose) were −5%, - 60%, −75% and −77% of mean baseline at dose levels of 0, 10, 30 and 100 mg/kg, respectively (Fig. 5E). Similar reductions in T_FH_ cells were also observed and the mean number of absolute T_FH_ counts at 28 days post-dose were+7%, −66%, −81%, and −86% of mean baseline values at dose levels of 0, 10, 30 and 100 mg/kg, respectively (Fig. 5F). The magnitude of the reduction in T_M_, and T_FH_ cells was therefore similar at dose levels of 30 and 100 mg/kg KY1044 did not appear to significantly affect the levels of circulating T_Reg_ (Supplementary Fig. 9C). However, this could be due to the fact that T_Regs_ are more difficult to quantify than T_M_ and T_FH_ cells since they only represent less than 3% of circulating CD4^+^ cells in monkeys (Fig. 5D). KY1044 did not notably affect CD8^+^ T cells in blood since the absolute counts were within the range observed during the pretreatment period (Supplementary Fig. 9D). As in the mouse study, we did not observe a significant decrease in ICOS^high^ cells such as T_M_or T_Reg_ or in ICOS^low^ CD8^+^ T cells in the in the lymph node (or spleen) of treated monkeys at termination the day after the last dose (Supplementary Fig. 9E-G). In summary, these data confirmed that KY1044 elicited a rapid and long-term depletion of ICOS^high^ cells in peripheral blood but not in lymphoid tissues of NHP’s.

### Increased IFN-γ and TNF-α expression in response to KY1044 *in vivo*

As presented above, KY1044 mIgG2a monotherapy modifies the intratumoural immune contexture with a significant decrease in T_Reg_ in the TME and an increase in the CD8 to T_Reg_ ratio. To demonstrate that KY1044 is also associated with activation of intratumoural T_Eff_ cells *in vivo*, we first assessed the expression of CD69 on the surface of CD8^+^ T cells. CD69 is a rapidly induced lymphoid marker used for the early detection of T cell activation (*38, 39*). We analysed T cells from CT26.WT tumours for CD69 expression, 24 hours after a second dose of KY1044 mIgG2a and compared this to the expression on T cells in the control group. As shown in Fig. 6A, KY1044 mIgG2a treatment was associated with an increase in CD69 expression, which was significant (P<0.05) at doses of 1 and 3 mg/kg. Similarly, the analysis of CD69/CD44 double positive cells also demonstrated a significant increase (p<0.05) of CD8 activation in response to KY1044 at a dose of 1mg/kg (Fig 6A).

**Figure 6:**
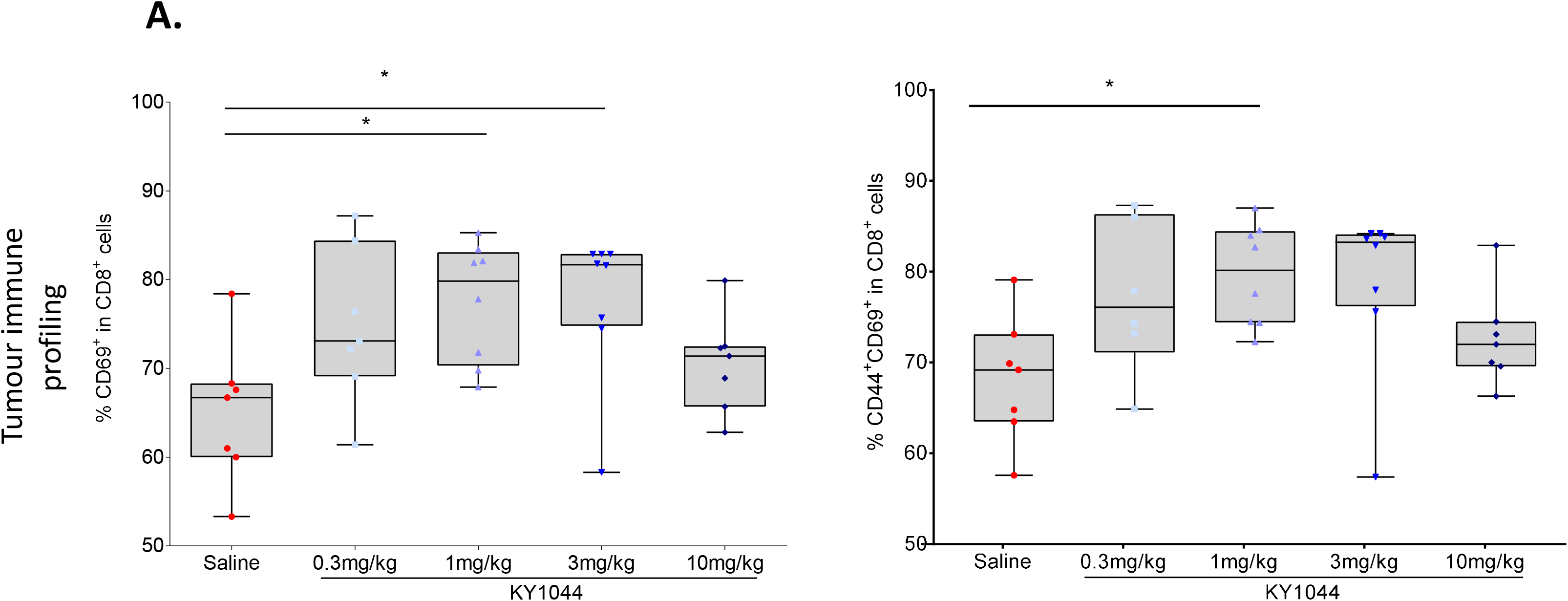
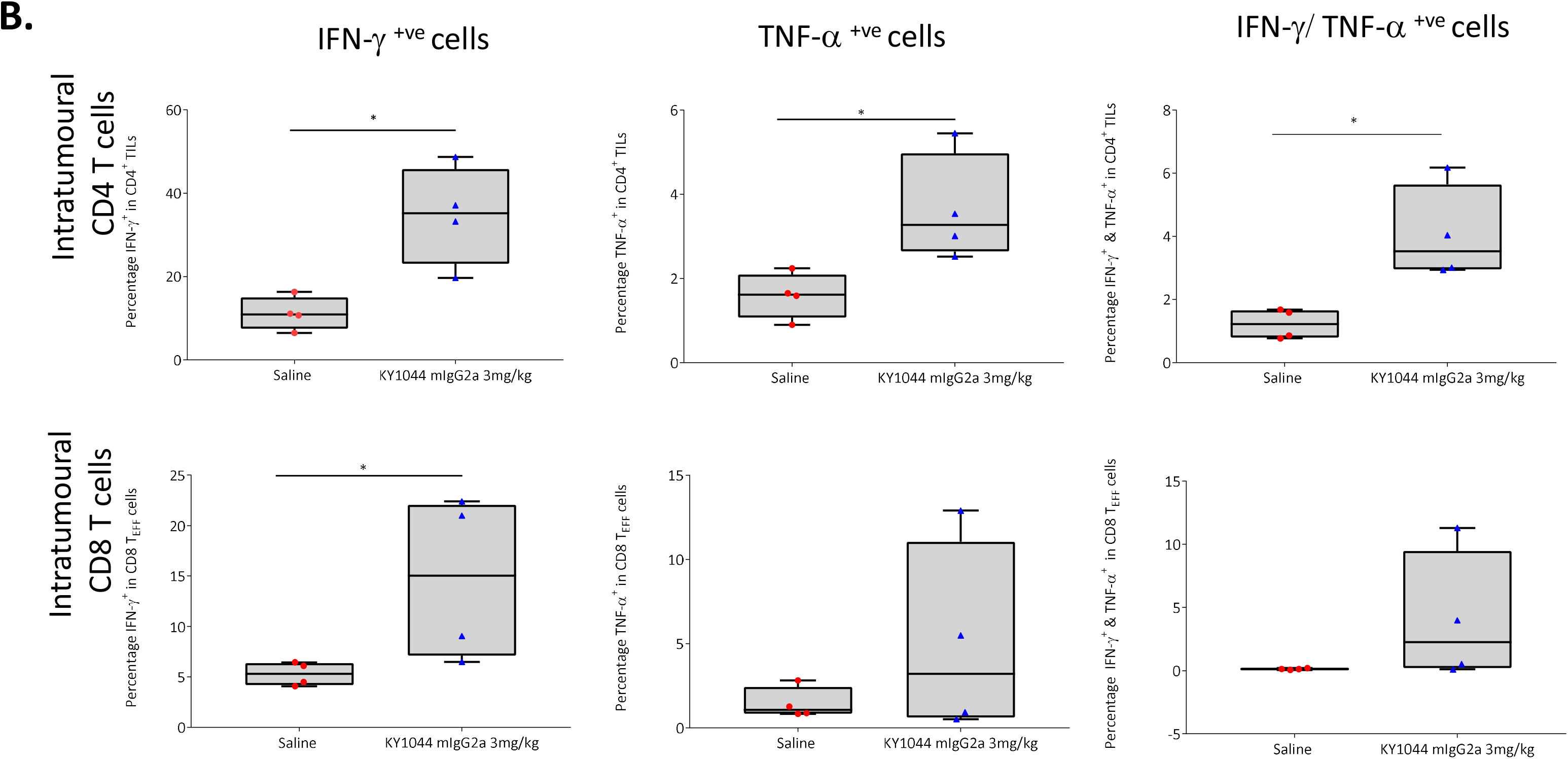
KY1044 mIgG2a activates intratumoural CD8 T cells as shown by the increase in CD69 and CD44 expression and increases the expression of Th1 cytokines IFN-γ and TNF-α in vivo. (A) KY1044 mIgG2a i.p injection results in an increase percentage of intratumoural CD8 T cells expressing the activation marker CD69 on their surface (n=7/8 mice per groups). Measurement of CD69 and CD69/CD44 double positive cells were performed 24 hours after a second dose of KY1044 mIgG2a (see figure 5A). (B) Graphs showing the flow cytometry analysis of Intracellular IFN-γ and/or TNF-α expression in intratumoural CD8+ and CD4+ T cells from CT26.WT tumour collected 7 days after i.p injection of saline or KY1044 mIgG2a at 3 mg/kg. Groups of mice (n = 4 mice/group). * p<0.05 Tukey’s multiple comparisons test

In a separate experiment, we also examined KY1044-dependent production of IFN-γ and TNF-α by CD4 and CD8 T cells. Both are crucial cytokines that play important roles in for the surveillance and inhibition of tumour growth *in vivo*(*40*). Using an intracellular staining approach on intratumoral CD4^+^ and CD8^+^ T cells collected 7 days post treatment (3 mg/kg of KY1044 mIgG2a), we demonstrated a significant increase in intracellular IFN-γ and/or TNF-α production in response to KY1044 mIgG2a, arguing for an improved anti-tumour phenotype of the intratumoural T cells (Fig. 6B). Although, one cannot differentiate between direct activation (through direct ICOS engagement on T_Eff_ cells) or indirect activation (via T_Regs_ depletion), the present immunophenotyping of intratumoural T cells following KY1044 mIgG2a dosing in vivo, revealed that the anti-ICOS was effectively associated with activation of both ICOS^Low^ CD8^+^ and CD4^+^ T_Eff_ cells

### Increased efficacy of KY1044 mIgG2a at an intermediate dose in combination with anti-PD-L1

The pharmacodynamic studies presented above confirmed that KY1044 mIgG2a effectively depletes ICOS^high^ cells such as T_Reg_ and result in activation of ICOS^Low^ T_Eff_ cells (albeit not sufficient to trigger anti-tumour efficacy as monotherapy in the CT26 model, Fig. 4A). However, we noticed that when used at high doses, KY1044 mIgG2a also affected the numbers of intratumoural CD8^+^ T cells (Supplementary Fig 8), resulting in a lower CD8^+^ T_Eff_ to T_Reg_ cell ratio. Since a higher baseline CD8^+^ T_Eff_ to T_Reg_ cell ratio has been shown to correlate positively with a response to immune checkpoint blockers such atezolizumab (*41*), we aimed to determine if the anti-tumour efficacy would be superior at an intermediate dose. For this, we repeated the CT26.WT efficacy study using a range of different doses of KY1044 mIgG2a (20, 60 and 200 μg/dose) combined with a fixed dose of anti-PD-L1 (200 μg/dose). As shown in Fig. 7, all the combination treatments were all associated with an improved response when compared with the saline or anti-PD-L1 monotherapy controls. Overall, this experiment demonstrated an improved response at the intermediate dose of 60 μg/dose of KY1044 mIgG2a, with 90% of the mice being tumour free and/or still on study by the study endpoint (day 54 post tumour cell implantation).

**Figure 7:**
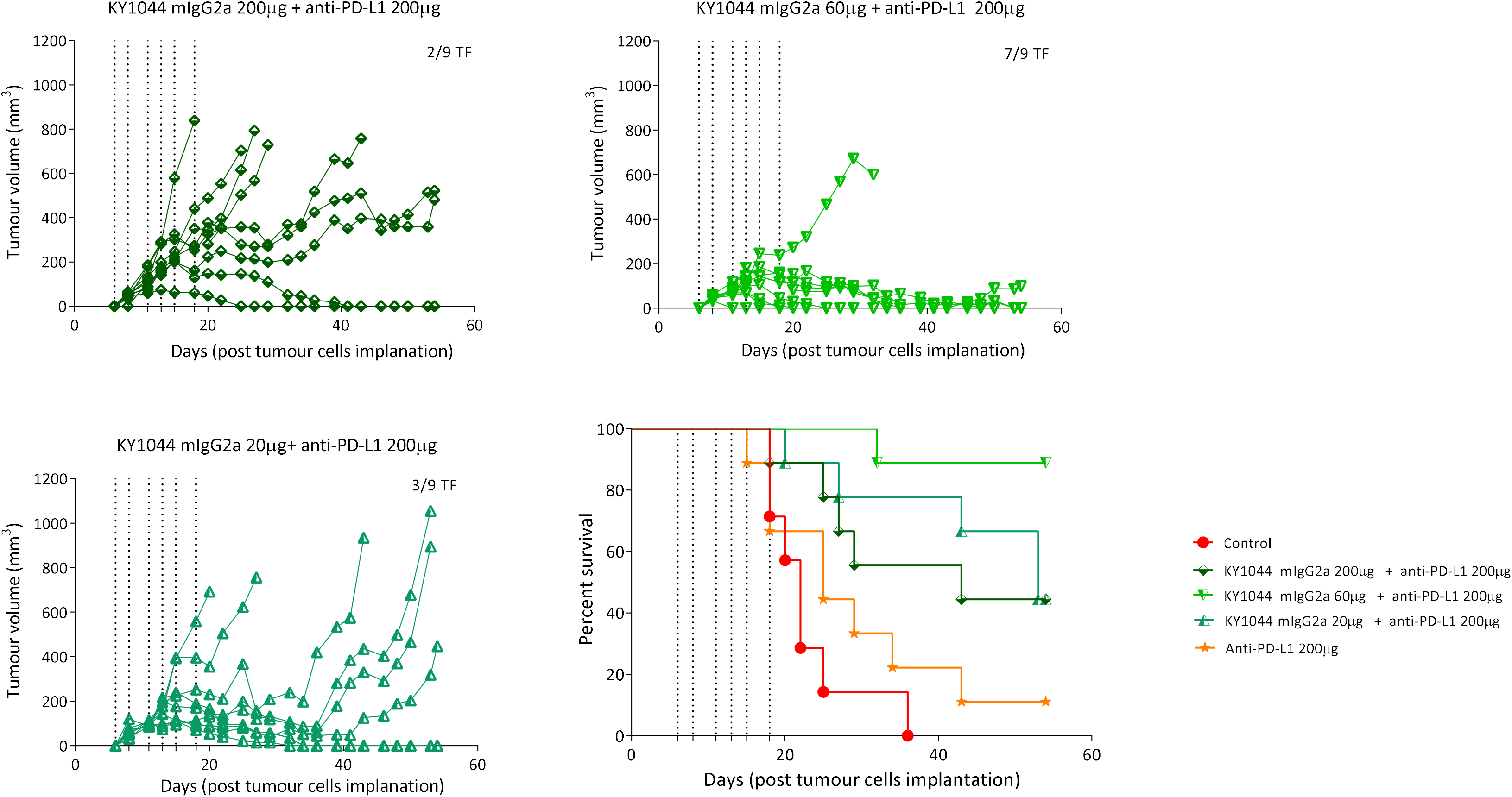
Better combination efficacy was obtained at intermediate dose of KY1044 mIgG2a. Spider plots showing individual mouse tumour volumes from BALB/c mice (n=9 per group) harbouring subcutaneous CT26.WT tumours and treated with different doses of KY1044 mIgG2a (20, 60 and 200 µg/dose) with a fixed dose of anti-PD-L1 (200 µg/dose). Kaplan-Meier plot depicting the survival of mice injected with CT26.WT-tumour and treated with different doses of KY1044 mIgG2a/anti-PD-L1 combination. The dosing days are indicated by vertical dotted lines.

## Discussion

The approval of ICIs as anti-cancer agents has been considered as a paradigm shift, providing a novel approach that adds to existing therapies used to fight cancer. However, responses to these checkpoint inhibitors are not universal, and many patients exhibit intrinsic or acquired resistance (*2*). Although the mechanisms of resistance are diverse, a strongly immunosuppressive TME can underlie the lack of response to ICIs in certain patients. Notably, intratumoural T_Reg_ are key cellular players involved in establishing and maintaining such an immunosuppressive TME. In addition, the high content of T_Reg_ is associated with poor prognosis for several tumour types and poor response to ICIs. For this reason, reducing T_Reg_ content and improving the ratio of effector T cells to T_Reg_ cells has emerged as an attractive strategy for improving the response to ICIs. This could be achieved via different strategies, including blocking the recruitment, function, expansion or survival of T_Reg_, and a number of molecules that could be targeted to modulate T_Reg_ are currently under investigation (*42*). These include CTLA-4, CD25, GITR, OX40, CCR8, CD137 and CCR2. However, the differential expression of these targets (i.e. between different immune cell subsets) is crucial to preferentially target T_Reg_, since none of these targets are specific to T_Reg_ (*8*). Moreover, in order not to affect the homeostasis of non-tumoural tissues and trigger severe organ-specific autoimmune diseases, an ideal target should be expressed at different levels on T_Reg_ in the TME and in other organs. In the present study, we performed extensive immunophenotyping of human, NHP and mouse samples at the mRNA and protein levels and demonstrated that ICOS expression differs between T cell subsets, with the highest expression observed on the surface of T_Reg_. In addition, using human and mouse tumour tissues, we confirmed that ICOS expression is higher in the TME than in the periphery (blood or spleen) or within the matching non-malignant tissue. Altogether our data suggests that due to its differential expression on different T cells subsets, ICOS represents a relevant and attractive target for an intratumoural T_Reg_ depletion strategy.

Another key aspect when developing novel anti-cancer therapies is to find the ‘right’ patients and the ‘right’ tumours. Our pre-clinical translational work has identified head and neck, gastric, oesophageal, lung, bladder, skin and cervical cancers as tumours with a high content of ICOS-positive cells. However, co-staining of ICOS with FOXP3 (used as a marker of T_Reg_) showed that these tumours differs by their incidence of ICOS^+^ T_Reg_. For example, lower ICOS^+^ T_Reg_ content was observed in gastric and bladder cancers, while cervical, oesophageal and head and neck cancers were more infiltrated by ICOS^+^ T_Reg_. Since the levels of ICOS expression on different cells subsets should affect the response to anti-ICOS therapies with a dual mode of action, our work suggests that the specific expression of ICOS on T_Reg_ may support a patient selection strategy. In addition, ICOS induction on T_Reg_ has been reported in chronic infectious disease (*43*). This is also relevant since several of the above preferred tumours that have high levels of ICOS^+^ T_Reg_ are also often associated with chronic viral or bacterial infections (*44*). At this stage, it is not yet clear if these T_Reg_ are pathogen-specific and are inhibiting the anti-tumour immune response as a “bystander effect”. However, one could hypothesise that the depletion of these cells should be beneficial for treating cancers associated with chronic viral and/or bacterial infections.

As discussed above, the expression of ICOS in tumours and the preferentially high levels of ICOS expression on intratumoral T_Reg_ highlight the potential of this protein for a depletion strategy through Fcγ receptor engagement on NK cells (ADCC) and macrophages (ADCP). Noteworthy, the fact that ICOS knockout is not embryonically lethal (unlike CTLA-4 knockout) also argues that the depletion of ICOS-positive cells would be better tolerated and not associated with severe autoimmune disease usually described with the targeting of CTLA-4 (*11, 45–47*). Using the Intelliselect^TM^ platform (*23, 24*), we identified and characterised a fully human anti-ICOS antibody called KY1044 that is being developed as an effector enabled human IgG1 antibody. We demonstrated *in vitro* that KY1044 has a dual mechanism of action, namely the potential to deplete ICOS^high^ cells via ADCC (through the engagement of FcgRIIIa) but also as a co-stimulatory pro-inflammatory molecule on cells expressing lower levels of ICOS, such as CD8^+^ T_Eff_ cells (through FcgR-dependent clustering). By reformatting KY1044 into a mouse IgG2a antibody, we also showed monotherapy efficacy in lymphoma models and combination efficacy (with anti-PD-L1) in models of solid tumours that are known to be resistant to PD1/PD-L1 blockade. Interestingly, in the CT26.WT tumour model, we showed better efficacy when combining anti-PD-L1 with KY1044 mIgG2a (effector enable) than with a poorly depleting KY1044 mIgG1 antibody. Similarly, in this model, we showed better anti-tumour efficacy when combining KY1044 mIgG2a with anti-PD-L1 than with anti-PD1. Although it is not clear why such a difference was observed in this model, one could postulate that anti-PD1, which blocks both PD-L1 and PD-L2, may be associated with a different phenotype to anti-PD-L1, which affects both PD1 and CD80 biology (*48*). Speculatively, there is also the possibility that both antibodies may interfere with each other’s functions within the immunological synapse, as both ICOS and PD1 are often expressed on the same cells and can affect each other pathways (*49–51*).

There is ongoing debate regarding the value of mouse models when predicting the depletion potential of effector function enabled antibodies targeting T_Reg_ in human (*8, 9, 22, 52, 53*). Experiments in mouse models have shown almost complete depletion of intratumoural CTLA-4 positive T_Reg_ whereas depletion of these cells in patients treated with ipilimumab remains disputed. The differences in FcgR expression and Fc/FcgR interaction on effector cells (NK and macrophages) between rodents and primates may have contributed to these discrepancies(*54*). Here, we performed pharmacodynamic studies in both mice (using mIgG2a) and NHPs (using hIgG1) to confirm that KY1044 could decrease the frequency of ICOS^high^ cells *in vivo* in both species. With the mouse work, we clearly demonstrated that KY1044 modifies the tumour immune contexture through reduction of T_Reg_ and by improving the CD8^+^ T_EFF_ to T_Reg_ cell ratio. Although this effect on ICOS-expressing cells was not sufficient to induce monotherapy efficacy in the CT26.WT tumour model, the change in immune cell contexture strongly sensitized the tumours to respond to anti-PD-L1 in a synergistic manner. Importantly, we demonstrated a decrease in ICOS^high^ cells in the blood of NHPs treated with KY1044 hIgG1. Depletion was also observed in single dose study in cynomolgus monkeys at dose levels as low as 0.1 mg/kg (data not shown). Together, these data in NHP suggest that KY1044 should be able to deplete ICOS^high^ cells in humans.

While the depletion of T_Reg_ is an attractive therapeutic strategy to re-activate host antitumor immunity, it is important that such an approach does not affect all ICOS^+^ T_Reg_ cells in all tissues as this could be associated with autoimmune side effects(*55, 56*). Here, we showed that, while T_reg_ were depleted in the tumour and the blood of treated animals, no significant depletion was observed in the spleen of treated mice or in the lymph node or spleen of treated NHPs. Although the reasons behind this selective depletion is not fully understood, this response has been shown for other depleting antibodies and, in our case, could be explained by the lower expression of ICOS on the surface of T cells in lymphoid tissues (supplementary Fig. 1). Altogether, these data suggest that KY1044 has the potential to deplete ICOS^high^ cells without affecting all cells expressing ICOS in the circulation and lymphoid tissues. This was also indirectly confirmed in the tumour rechallenge experiment, in which a long-term immune memory response was observed after treatment with KY1044 mIgG2a. Since memory T cells are known to express ICOS, this experiment suggests that that KY1044 mIgG2a does not kill all ICOS^+^ T cells. Finally, we also demonstrated that KY1044 has co-stimulatory properties, as shown by the activation and secretion of cytokines such as IFN-γ and TNF-α both *in vitro* and *in vivo*. Although no effect on T cell proliferation were observed, we noted a strong effect on cell morphology in response to KY1044-dependent ICOS stimulation.This co-stimulatory agonistic property was shown to be dependent on clustering of the antibody and on the concomitant engagement of the T cell receptor.

The depletion of ICOS-expressing cells other than T_Reg_ in the blood of cynomolgus monkeys was not associated with any adverse toxic effects and the depletion of up to about 80% of the T_M_ cells in peripheral blood is unlikely to present a significant risk of loss of immunologic memory to previous pathogenic antigen exposure or vaccinations. In humans, circulating T cells, of which memory cells represent about 40-60% in adults, represent <3% of total T cells in the body (*57*). T_M_ cells in peripheral blood only have a lifespan of about 5 months (*58*) and are replenished from a large pool of antigen-specific memory cells which reside in lymphoid tissues (mucosal sites, skin, spleen, lymph nodes and bone marrow; (*57*)) and KY1044 did not deplete T_M_ cells in lymph nodes or spleen of NHP.

Finally, while assessing the effect of KY1044 on the CD8^+^ T_EFF_ to T_Reg_ intratumoural ratio, we noticed that, although the treatment improved the ratio at all doses tested, a bell shape response pattern was observed, with a lower ratio obtained at the lowest and highest concentrations of KY1044 (0.3 and 10 mg/kg, respectively). At the highest dose, this could be explained by the reducing trend (albeit not significant) in the numbers of intratumoural CD8^+^ T cells. Although the cause of this decrease is not clear, there is a possibility that either cells expressing low levels of ICOS can be depleted at high doses of KY1044 or that the activation of T_Eff_ cells in response to KY1044 is associated with increased ICOS expression, which could make these cells more sensitive to depletion. It was also noted that, while all combinations of anti-PD-L1 with different doses of KY1044 mIgG2a were shown to trigger an anti-tumour immune response, the intermediate dose of 60 µg of KY1044 resulted in the strongest response. Finding the most effective dose in patients will therefore require a thorough monitoring of immune cell contents in the blood and TME.

Altogether, our tumour efficacy and pharmacodynamic studies clearly demonstrate that KY1044 is pharmacologically active, modifies the intratumoural immune contexture and induces a strong and long-lasting anti-tumour immune response. These findings, therefore, warrant the further assessment of KY1044 – as a monotherapy or in combination with anti-PD-L1 as a potential treatment for solid tumours.

## Material and Methods

### Cell lines used in the study

MC38 cells were obtained from National Institute of Cancer under a license agreement. J558 (ATCC® TIB-6™), CT26.WT (ATCC® CRL-2638™), A20 [A-20] (ATCC® TIB-208™), EL4 (ATCC® TIB-39™), B16-F10 (ATCC® CRL-6475™) and MJ [G11] (ATCC® CRL-8294™) cell lines were obtained from the ATCC. The specific pathogen free status of these cells was confirmed by PCR screening for mouse/rat comprehensive panel (Charles River). MC38, J558 and B16-F10 cells were cultured in antibiotic free DMEM (Gibco, 41966-029) + 10% FBS (Gibco, 10270) complete cell culture media. CT26.WT cells were cultured in antibiotic free RMPI (Gibco, 2187) + 10% FBS complete cell culture media. A20 cells were cultured in antibiotic free RMPI (Gibco, 2187) + 10% FBS + 0.05mM 2-mercaptoethanol (Gibco, 21985-023) complete cell culture media. MJ cells were culture in IMDM + 20% FBS (I20 media). The passage number of tumour cells were kept below 10 generations.

### Gene expression analysis

#### TCGA data analysis

The standardized, normalised and batch corrected RNA sequencing data collected as part of the TCGA consortium was downloaded from the UCSC Xena platform (*59*). Samples classified as non-tumour tissue were excluded from the dataset. Single sample gene set enrichment analysis (ssGSEA (*60*)) was performed for a gene set consisting of Icos and Foxp3 using the R package GSVA (*61*). Samples were grouped by primary disease and the ssGSEA scores for each group were compared across the primary disease groups.

#### Single-cell RNA sequencing

PBMC and tumour cell suspensions from 5 NSCLC donors were processed using the BD Rhapsody single cell analysis system. Sequencing libraries were generated using the Immune Response Human targeted panel and sequenced using a 2 × 75 bp paired-end run on the Illumina HiSeq 4000 System. Reads were processed by applying the BD Rhapsody processing pipeline to generate cell count matrices. The counts were then filtered, normalised and visualised using R and Bioconductor packages for single-cell RNA-seq data (*62–65*). Cell type specific gene sets were constructed by performing a literature search and cells were classified into one of 27 cell types (Supplementary Table 1) using the R package AUCell (*60*)

### Flow Cytometry

#### Mouse Studies

Tumours and spleens were harvested from CT26.WT tumour bearing mice at day 16 post tumour cell implantation. Tumours were dissociated into single cell suspensions using Miltenyi Tumour Dissociation Kit for mouse tissues (Catalogue no. 130-096-730) as per the manufacturer’s instructions. Spleens were placed into C-tubes (Miltenyi Biotec, Catalogue no. 130-096-334) dissociated into single cell suspensions using gentle MACS^TM^ dissociator. Tumour samples were filtered through 70 µm nylon filters (Falcon, Catalogue no. 352350). Spleens samples were filtered through 40 µm nylon filters. Red blood cells in the spleens were lysed using RBC lysis buffer (Sigma, Catalogue no. R7757). For flow cytometry profiling, 2 x 10^6^ tumour samples or 1 x 10^6^ spleen samples were plated in 96 well plate (Sigma, Catalogue no. CLS3957). Cell suspensions were pre-incubated with Live/Dead™ fixable Yellow Dead Cell Stain Kit (Invitrogen), according to the manufacturer’s instructions. Prior to antibody labelling cell were incubated with Fc receptor blocking solution (Anti-CD16/CD32) at 4°C for 10 minutes. Cell surface staining was performed at 4°C for 30 minutes using fluorochrome-conjugated anti-mouse antibodies. Intracellular and intranuclear staining was performed using Foxp3 staining buffers (ThermoFisher Scientific, catalogue: 00-5523-00) according to manufacturer’s instructions. All flow cytometry antibodies or isotype controls were purchased from ThermoFisher Scientific. Antibodies used include: anti-CD45 (30-F11), anti-CD3 (17A2), anti-CD4 (RM4-5), anti-CD8 (53-6.7), anti-CD25 (PC61.5), anti-ICOS (7E.17G9), anti-Foxp3 (FJK-16s), anti-CD69 (H1.2F3).

For the tumour infiltrating T cell cytokine staining, single cell suspensions from CT26.WT tumours were plated at 1 x 10^6^ cells per well in RPMI + 10% FBS cell culture media with 1 x Brefeldin A solution (eBioscience, catalogue no: 00-4506-51) for four hours in a cell culture incubator at 37°C 5% CO2. Cells were surface stained as above and subsequently fixed/permeabilised for intracellular staining with anti-IFN-γ (XMG1.2) and anti-TNF-α (MP6-XT22). All flow cytometry data was acquired using Attune FxT flow cytometer and data was analysed using FlowJo^TM^ software V10.

#### Monkey study

Blood samples (0.1 mL) were collected into tubes containing sodium EDTA or lithium heparin, mixed and red blood cells (RBCs) were lysed with lysis buffer containing ammonium chloride. The remaining leukocytes were washed with FACS buffer (PBS with 2% foetal bovine serum), stained with antibody cocktail by incubation in the dark for 45 minutes at room temperature and washed. Tissue samples (lymph node), taken from animals at termination were mechanically disrupted (Medimachine, Becton Dickinson GmbH, Heidelberg, Germay (BD)), single cell filtered and stained as for blood. Intracellular staining (for FoxP3) was carried out by permeabilization with 10x FACS lysing solution (1:5 with Aqua dest. +0.1% Tween-20) prior to incubation in the dark at room temperature for 45 minutes and wash steps. Antibody cocktails (including appropriate isotype antibodies) were prepared on the day of use and stored in the dark.

All flow cytometry antibodies or isotype controls were purchased from BD, eBioscience (via Fisher Scientific GmbH, Schwerte, Germany) or BioLegend (Koblenz, Germany). Antibodies used included: anti-CD3 (SP34), anti-CD20 (L27 or 2H7), anti-CD14 (M5E2), anti-CD4 (M-T477), anti-CD8 (SK1), anti-CD28 (CD28.2), anti-CD25 (M-A251), anti-Foxp3 (PCH101), anti-CD95 (DX2) and anti-CD185/CXCR5 (MU5UBEE). After staining, cells were washed and fixed in 1 × BD Stabilizing Fixative. Lymphocytes were gated by forward scatter (FSC), sideward scatter (SSC) and CD45. Multicolour flow cytometric analysis was performed using the following leukocyte phenotypic characteristics: CD4^+^ T helper cells: CD3^+^ /CD4^+^ /CD8^-^ /CD14^-^ /CD20^-^; CD8+ cytotoxic T cells: CD3^+^/CD4^-^/CD8^+^/CD14^-^/CD20^-^; Total memory CD4 T cells: CD28^+/DIM/-^ CD95^+high^; Follicular T helper cells: (CD4^+^/CD185^+^) and Regulatory CD4 T cells: CD25^+^/ FoxP3+. Acquisition of flow data was performed on a FACSVerse^TM^ flow cytometer (BD) and relative percentages of each of these subpopulations were determined using FlowJo^TM^ software. Fifty thousand events were counted for all analyses. The absolute numbers of each of the subpopulations were determined for blood samples by calculations based on haemoanalyser analysis of whole blood.

#### Human studies

NSCLC tissue and whole blood from the same patients was obtained from consented subjects, with ethical approval for analysis of protein, RNA and DNA content. The NSCLC tumour samples were dissociated into single cell suspensions using enzymatic digestion, whilst whole blood was processed into PBMCs using density centrifugation, all specimens were cryopreserved in Gibco Recovery™ cell culture freezing medium before used. Alongside NSCLC specimens, whole blood samples from 5 healthy individuals were collected and processed into PBMCs by density centrifugation. Flow cytometry was performed after cells were thawed and incubated in RPMI-1640 + 10% FBS supplemented with 100 units of RNase free DNase at 37°C 5% CO2 for 30 minutes. Cell suspensions were pre-incubated with Live/Dead™ fixable Yellow Dead Cell Stain Kit (Invitrogen) and Fc receptor blocking solution (Biolegend), before staining with fluorochrome-conjugated anti-human antibodies specific to anti-CD3 (UCHT1)/anti-CD45 (2D1), anti-CD4 (2A3), anti-CD8 (RPA-T8), anti-CD25(MA251), anti-CD45RA (H100), anti-ICOS (C398.4A). Cells were incubated for 1 hour at 4°C, washed and fixed overnight at 4°C with eBioscience™ intracellular fixation and permeabilization buffer. Anti-FoxP3 (236A-E7) staining was then performed in permeabilization buffer for 1 hour at 4°C before washing and resuspending cells in DPBS (Gibco). All flow cytometry data was acquired using Attune FxT flow cytometer and data was analysed using FlowJo^TM^ software V10.

### Immunofluorescence and digital pathology

An IHC protocol using a semi-automated method on the Ventana Discovery Ultra platform (Roche) was developed for co-staining of ICOS and FoxP3. Method was optimised in FFPE tonsil tissue, using clone D1K2T for the detection of ICOS (Cell Signaling), while clone 236A/E7 (Abcam) was chosen for FoxP3 detection. Methods were applied for the staining of tumour microarrays (TMA) from cervical (CR2088), oesophageal (ES2082), lung (LUC2281), head and neck (HN802a), gastric (STC2281) and bladder (BL2081a) cancer patients. The TMAs (tissue microarrays) were sourced from US Biomax, Inc. All stained slides were scanned at a magnification of 20x equivalent magnification, generating images which were quantified using the Indica Labs Halo platform to determine the number of ICOS/FoxP3 single and double positive cells per mm^2^.

### In vitro ADCC assays

The ADCC activity of KY1044 human IgG1 (produced at Kymab) was first tested in vitro using an ADCC reporter bioassay (Promega) according to the manufacturer’s instructions. In brief, CHO cells expressing either human, mouse, rat or cynomolgus ICOS as target cells were co-incubated with ADCC reporter cells expressing human FcγRIIIa (V158; Promega) at a 5:1 ratio. Serial dilutions of KY1044 or isotype control were added to the culture plates, incubated at 37 °C overnight and the luciferase activity was measured using the Bio-Glo Luciferase Assay System (Promega) on the EnVision Multilabel Plate Reader (Perkin Elmer). Graph data were normalised to background and plotted versus Log10 antibody concentration.

For the primary cells ADCC assay, human NK cells were purified from PBMC isolated from whole blood by density gradient centrifugation. NK cells were subsequently purified from the by immunomagnetic negative selection using the NK Cell Isolation Kit (Stemcell) as per manufacturer instructions. ICOS-transfected CCRF-CEM (ICOS CEM, target cells) were preloaded with the fluorescence enhancing ligand (BATDA) for 30 minutes in the dark at 37 °C. At various occasions the cells were loaded and/or washed in the presence of an inhibitor of organic anion transporters (1mM Probenecid) to avoid spontaneous dye release from cells. KY1044 was diluted (1:4 dilutions, 10 points, starting from 33.3 nM) in assay buffer. The target ICOS CEM cells (50µl/well), effector cell (50µl/well) and reagent dilutions (50µl/well) were co-cultured with 50µl of the diluted antibody at 37 °C, 5% CO2 for 2-4hrs. The effector NK cells:target cell ratio was fixed at a 5 : 1 ratio. A digitonin-based lysis buffer (Perkin Elmer) was prepared in parallel and used to determine complete target cell lysis (100%).

### In vitro agonism assays

For the MJ cells assays, KY1044 human IgG1 was presented to the MJ cells in 3 different formats: plate-bound, soluble or soluble plus F(ab’)₂ Fragments (Fc linker, 109-006-170, Jackson Immuno Research). For the plate bound assay, KY1044 and the hIgG1 isotype control were diluted 1:2 in PBS to give final antibody concentrations ranging from 10 μg/mL to 40ng/mL (10 points). 100 μL of diluted antibodies were coated in triplicate into a 96-well, high-binding, flat-bottom plate (Corning EIA/RIA plate) overnight at 4 °C and then washed. MJ cells (15,000 cells/well) were added to Ab-containing wells. For the soluble/cross-linked experiment: KY1044 and the isotype control were serially diluted 1:2 in I20 media (soluble Ab) or in I20 media containing 30 μg/mL of F(ab’)₂ Fragments (cross-linked Ab) to give an 2X Ab stock concentrations ranging from 20 μg/mL to 80 ng/mL (10 points). 50 μL of diluted Abs were added to 96-well with 50 μl of MJ cells (3×10^5^/mL). For both assays the cells were cultured for 72 hrs at 37 °C and 5% CO2 and Cell free supernatants were then collected and used to perform IFN-γ ELISA using the Human IFN-γ DuoSet assay (R&D system, DY285).

For the primary T cell assays, leukocyte cones were obtained (HTA IRAS project number 100345). PBMCs were isolated from whole blood by density gradient centrifugation. T lymphocytes were them purified using the Stemcell EasySep Isolation kit (cat#17951) following manufacturer’s instructions. In order to induce ICOS expression, isolated T-cells were cultured at 2×10^6^/mL in R10 media (RPMI 10% heat inactivated FBS) in the presence of 20 μl/ml of Dynabeads Human T-Activator CD3/CD28 (from Life Technologies). These activated primary T cells were tested as for the MJ cells assays in 3 different formats: plate-bound (5 μg/mL), soluble (15 μg/mL) or soluble plus F(ab’)₂ Fragments cross-linker. T-cell suspension were added to Ab-containing plates to give a final cell concentration of 1×10^6^ cells/ml and cultured for 72 hrs at 37°C and 5% CO2 until IFN-γ ELISA quantification. For the 3-step culture (stimulation-rest-costimulation assay), the T-cells were pre-stimulated by Dynabeads for 3-days to induce ICOS before being rested for 3-days to reduce their activation levels. These stimulated/rested T-cells were then cultured with KY1044 in the presence or absence of an anti-CD3 Ab (clone UCHT1, eBioscience) to assess the requirement of TCR engagement. The effect of ICOS co-stimulation was assessed after 72 hrs by measuring the levels of IFN-γ and TNF-α present in the culture (MSD multiplex assay).

For the gene expression analysis, T cells were harvested from the 3-step culture (stimulation-rest-costimulation assay) after 6-,hrs of plate-bound antibody stimulation. Total RNA was extracted from the cell pellets with the RNeasy Micro Kit (QIAGEN), quality controlled on the Agilent 2100 Bioanalyzer (Agilent Technologies) and subjected to SE50 sequencing following mRNA enrichment (BGI). The sequence reads were aligned using kallisto (*67*) and further processed using limma (*68*) and metascape (*69*) and GSVA (*61*).

### Mice

All mice for *in vivo* work was cleared through local ethical committee and was performed under Home Office license. 8 to 10 week-old wild-type female Balb/C or C57BL/6J mice were sourced from Charles River UK Ltd (Margate, UK) and housed in transparent plastic cages with wire covers (391 W x 199 L x 160 H mm, floor area: 500 cm^2^) containing Grade 6 Wood Chip which can be replaced with Lignocell (IPS Product Supplies Ltd, BCM IPS Ltd. London. WC1N 3XX) and bale shredded nesting material (IPS Product Supplies Ltd, BCM IPS Ltd. London. WC1N 3XX). Four to five Mice were housed per cage in a room with a constant temperature (19-23°C) and humidity (40-70 %) and a 12-hour light-dark cycle (lighting from 7 A.M. to 7 P.M.). Mice were provided with pellet food (CRM(P), Specialist Diet Services, Witham, UK) and RO water *ad libitum* using an automatic watering system.

### Tumour cell implantation and tumour measurement

Early passage (below P10) MC38 (3×10^6^ cells), J558 (1×10^6^ cells) CT26.WT (1×10^5^ cells), A20 (5×10^6^ cells), EL4 (1×10^4^ cells) and B16-F10 (1×10^5^ cells) tumour cells were prepared in either PBS or Matrigel (Corning, 354230) and the resulting cell suspension were injected subcutaneously into the flank of the mice (Study day 0). Prior to tumour cell implantation, mice were anaesthetised with isoflurane and the right flank of the mice was shaved. For the implantation, 100 µl of the cell suspension were injected using 25 G needles (BD Microlance TM 3. VWR 613-4952). Cell numbers and viability (required to be above 90%) were determined pre-implantation by the trypan blue assay. Tumour growth was measured using digital callipers three times a week until end of the study. The tumour volumes (mm^3^) was estimated using a standard formula: (L x W^2^) /2 (with L being the larger diameter, and W the smaller diameter of the tumour. All data were plotted using GraphPad Prism V10.

### Antibody dosing

For the *in vivo* tumour efficacy studies, both KY1044 mIgG2 and anti-PD-L1 (AbW) mIgG2a were produced by Kymab ltd. The anti-PD1 (clone RMP1-14) monoclonal antibody was purchased from BioXcell. Antibodies were dosed between 0.1 and 10 mg/kg or by flat dose of 20 to 200 μg/dose via the intraperitoneally route. For the efficacy and pharmacodynamic studies the treatment groups were not blinded. The *in vivo* depletion of CD8^+^ and CD4^+^ T cells were conducted using a flat dose 200 µg of anti-CD8a (53-6.7) and/or anti-CD4 (GK1.5). Dosing was performed twice a week for 3 weeks starting from day 3 post tumour cell implantation. The efficiency of the antibody mediated CD8 and CD4 T cell depletion was determined by flow cytometry of tumour, spleen, tumour draining lymph node (inguinal lymph node) and blood cells using an anti-CD3 antibody (17A2).

### Cynomolgus monkey Study

The pharmacodynamic effects of KY1044 in non-human primates were studied as part of a repeat dose toxicity study. Naïve male cynomolgus monkeys (Macaca fascicularis) were obtained from a certified supplier, group housed, allowed access to water ad libitum and fed on a pelleted diet for monkeys supplemented with fresh fruit and biscuits. Animals ranged from 4 to 7 years old and weighed 3-6 kg at time of dosing. Four groups of three cynomolgus monkeys received weekly intravenous (i.v.) doses of KY1044 (slow bolus over approximately 1 minute) for a month (5 doses in total) at dose levels of 0 (vehicle control; phosphate-buffered saline pH 7.4), 10, 30 or 100 mg/kg at a dose volume of 2 mL/kg. Blood samples were taken at 1 or 2 timepoints prior to dosing and at multiple timepoints up to 29 days after the first dose for measurement of serum KY1044 using a qualified ELISA assay, ICOS occupancy on CD4+ cells in blood determined using a validated flow cytometry method and/or immunophenotyping of whole-blood (described above). Scheduled necropsies were conducted 1 day after the final dose (Day 30) and spleen and mesenteric lymph nodes were taken for immunophenotyping.

### Statistical analyses

Unless otherwise stated in the Figures legend, the efficacy observed for the different treatment groups were compared using either T-test or one-way analysis of variance (ANOVA) and Tukey’s multiple-comparison post-hoc test. Differences between groups were significant at a *P* value of <0.05. Statistical analyses were performed with GraphPad Prism 10.0 (GraphPad Software, Inc., San Diego, CA).

## Supporting information

MJ cells treated with isotype control

MJ cells treated with anti-ICOS

## Acknowledgements

The authors are extremely grateful to Kymab’s molecular biology and antibody expression team as well as the Kymab BSU team. The authors would also like to thank Propath UK (Hereford, UK) and OracleBio (Biocity, UK) for assistance with IHC staining and digital image analysis as well as Stephanie Grote-Wessels (Covance Preclinical Services GmbH) for their support with this study. We also thank the US. National Cancer Institute’s Division of Cancer Treatment and Diagnosis, Developmental Therapeutics Program, Biological Testing Branch, Tumor Repository for providing the MC38 syngeneic tumor cell line.

## Author contributions

RCAS, AKT, MK, SH, JaC, EO, SQ, MMc for the study conception and design; AKT, GB, MK, NP, RR, RK ML, JC, DM,SoL, SB, LG, HA, HC,VW, QL, JoC, IK TM, ET, EP, JaC for the acquisition of data; RCAS, AKT, MK, GB, NP, SH, JoC, MMc, JaC, IK, VG Analysis and interpretation of data and RCAS, AKT, MK, SH Drafting of manuscript: EO, VG, SQ and MMc for the critical revision.

## Supplementary Materials

### Materials and Methods

#### Luciferase assays

Luciferase reporter assays: Jurkat-Lucia™ NFAT cells (Invivogen) stably expressing luciferase under the control of NFAT response elements were further transfected with either the human *Icos* gene sequence or a chimeric construct of *Icos* fused with *CD247* (CD3ζ). Following selection of stable transgene integration, a luciferase activity bioassay was performed. High binding 96-well assay plates were coated overnight with either anti-CD3 (UCHT-1, BD Biosciences; 10 µg/ml), anti-ICOS (C398.4A, BioLegend; 10 µg/ml), isotype control (HTK888, BioLegend; 10 µg/ml), anti-CD3 + anti-ICOS (5 µg/ml each) or anti-CD3 + isotype control (5 µg/ml each). Transgene expressing cells or control cells (untransfected Jurkat-Lucia™ NFAT cells) were seeded at 50,000 cells / well. Following overnight incubation luciferase activity was measured by adding BioGlo reagent (Promega) and reading luminescence on a plate reader. Luciferase activity was normalised and scaled to 100% by comparing to cells stimulated using Cell Stimulation Cocktail (eBioscience).

#### NHP ethics statement

The cynomolgus monkey study was conducted at Covance preclinical Services GmbH, Muenster, Germany in strict accordance with a study plan reviewed and approved by the local Institutional Animal Care and Use Committee (IACUC) of the testing facility and the German Animal Welfare Act. The study was performed according to DIRECTIVE 2010/63/EU OF THE EUROPEAN PARLIAMENT AND OF THE COUNCIL of 22 September 2010 on the protection of animals used for scientific purposes and the Commission Recommendation 2007/526/EC on guidelines for the accommodation and care of animals used for experimental and other scientific purposes (Appendix A of Convention ETS 123). The study was in compliance with the Kymab Working Practice on Experiments involving Animals and was approved by the Kymab Ethics Committee.

## Supplementary figure legends

**Supplementary figure 1:**
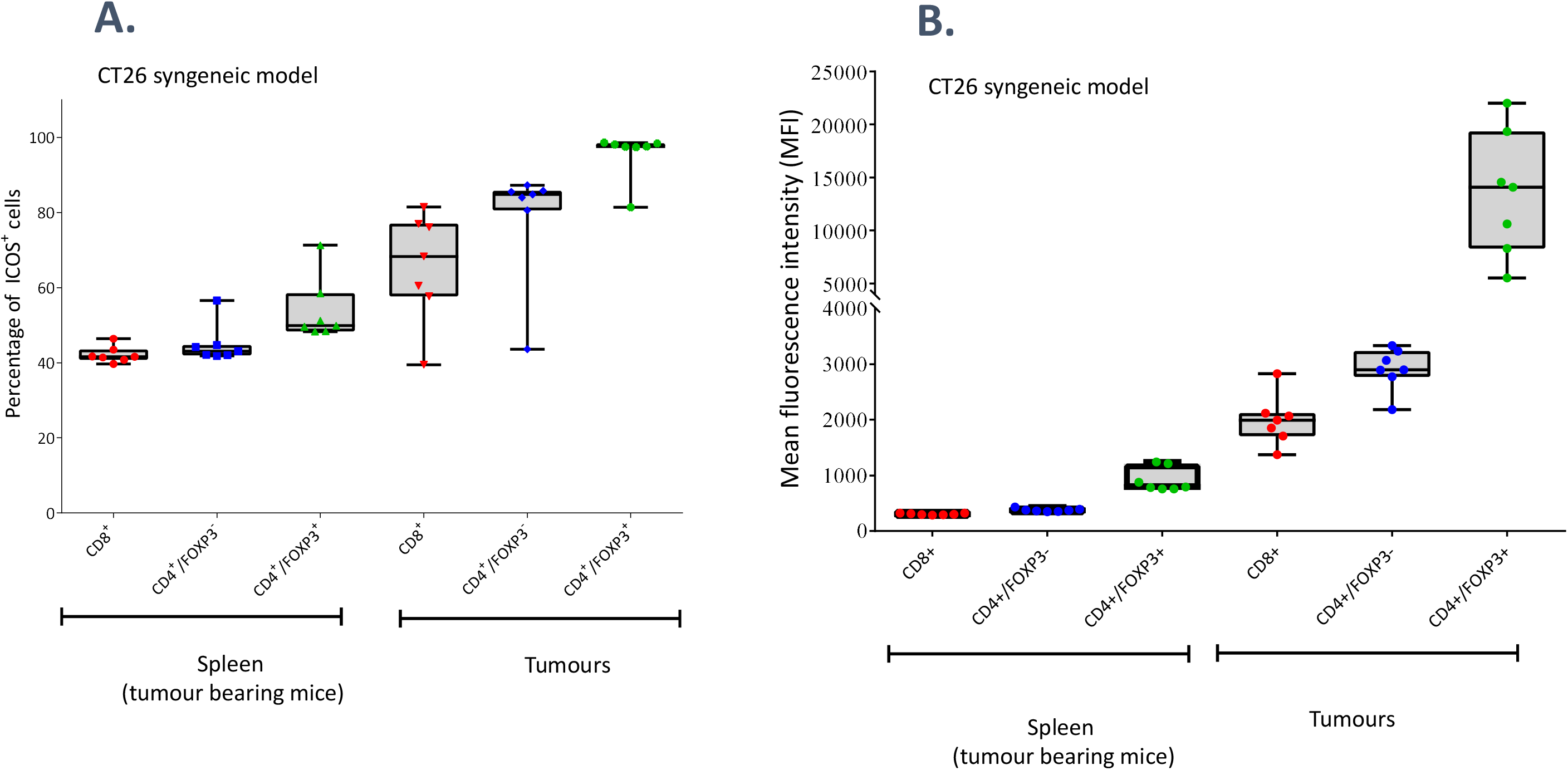
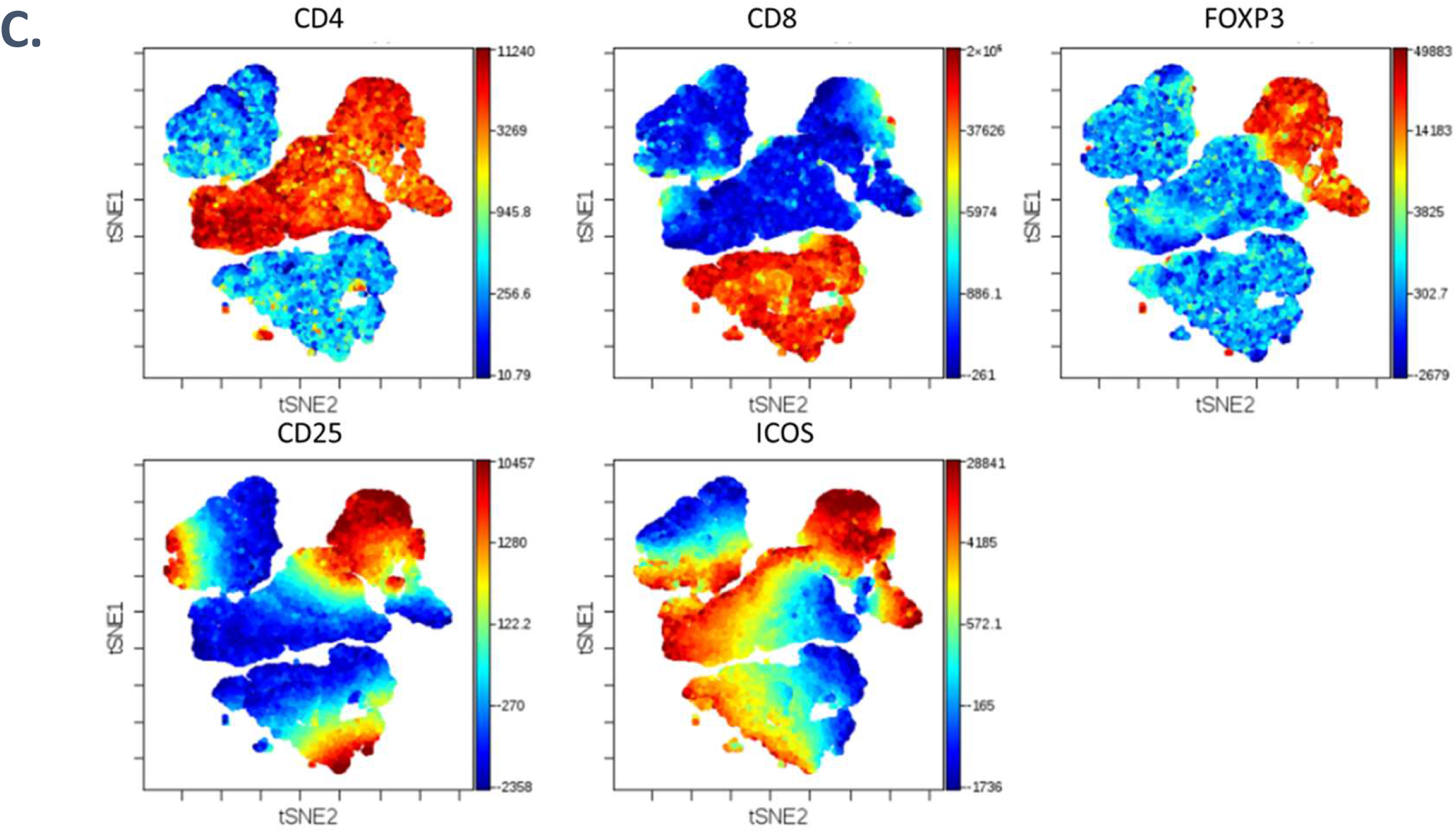
FACS analysis showing the (A) percentage of ICOS positive cells and (B) the relative ICOS expression in T cells derived for the spleen and tumours of CT26.WT tumour bearing mice. (C)) viSNE (visualization of t-distributed stochastic neighbor embedding) map of the T cell subpopulations (CD3+ cells) from untreated CT26.WT tumour samples (Cytobank). The colour scales depict the expression levels of selected markers (CD4, CD8, FOXP3, CD25 and ICOS) ranging from blue (low expression) to red (high expression) from the total T cell population that was organised in 2D based on similarity of expression.

**Supplementary figure 2:**
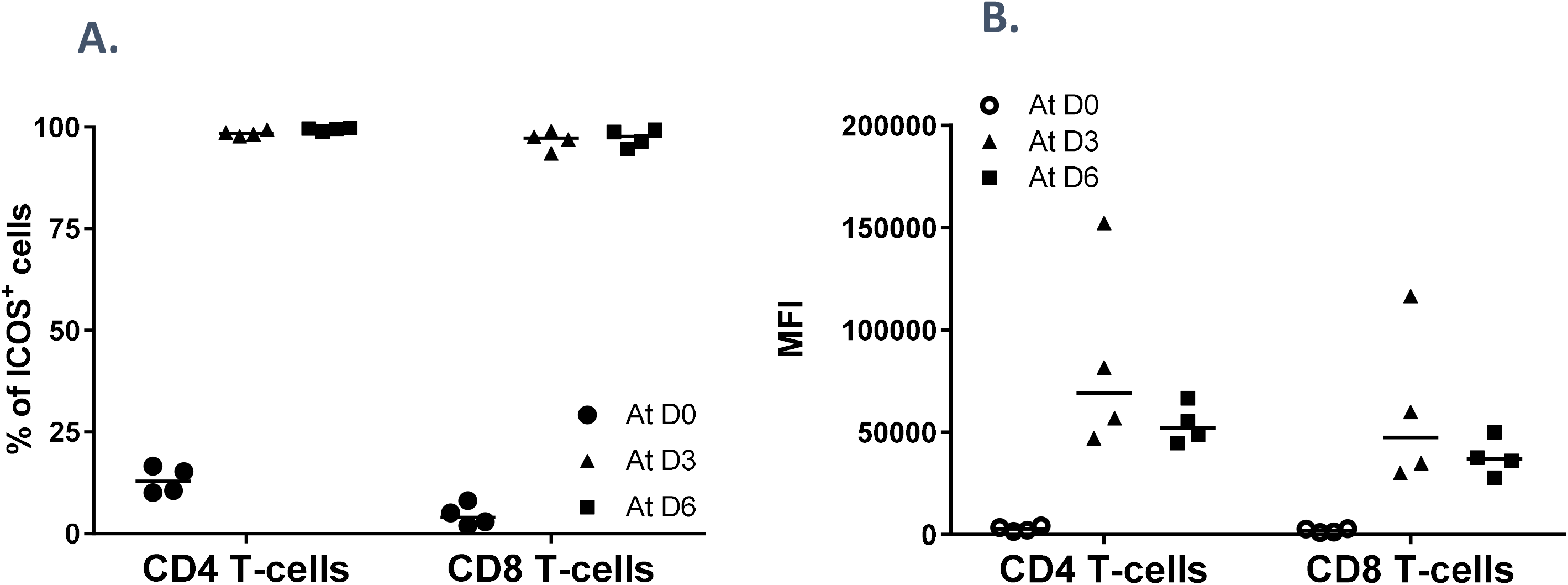
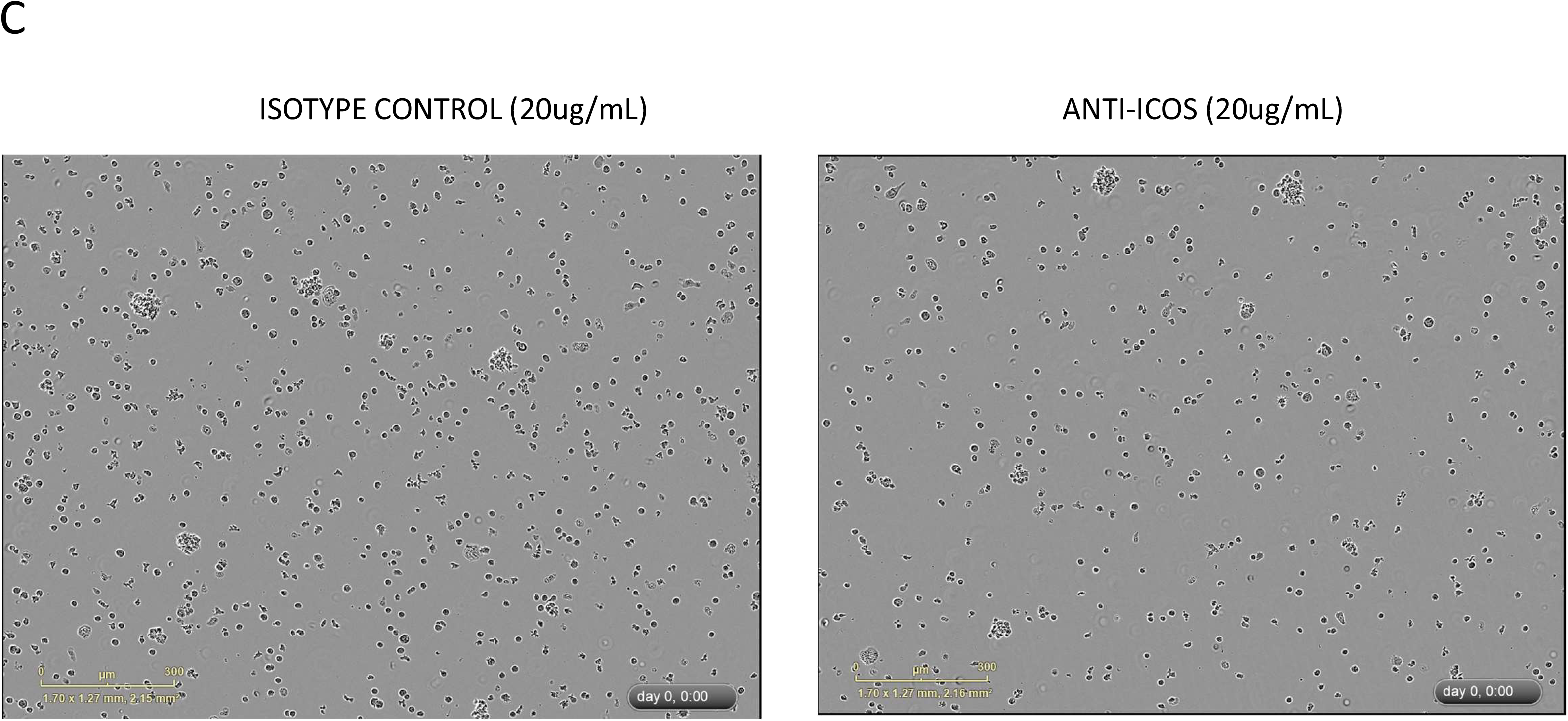
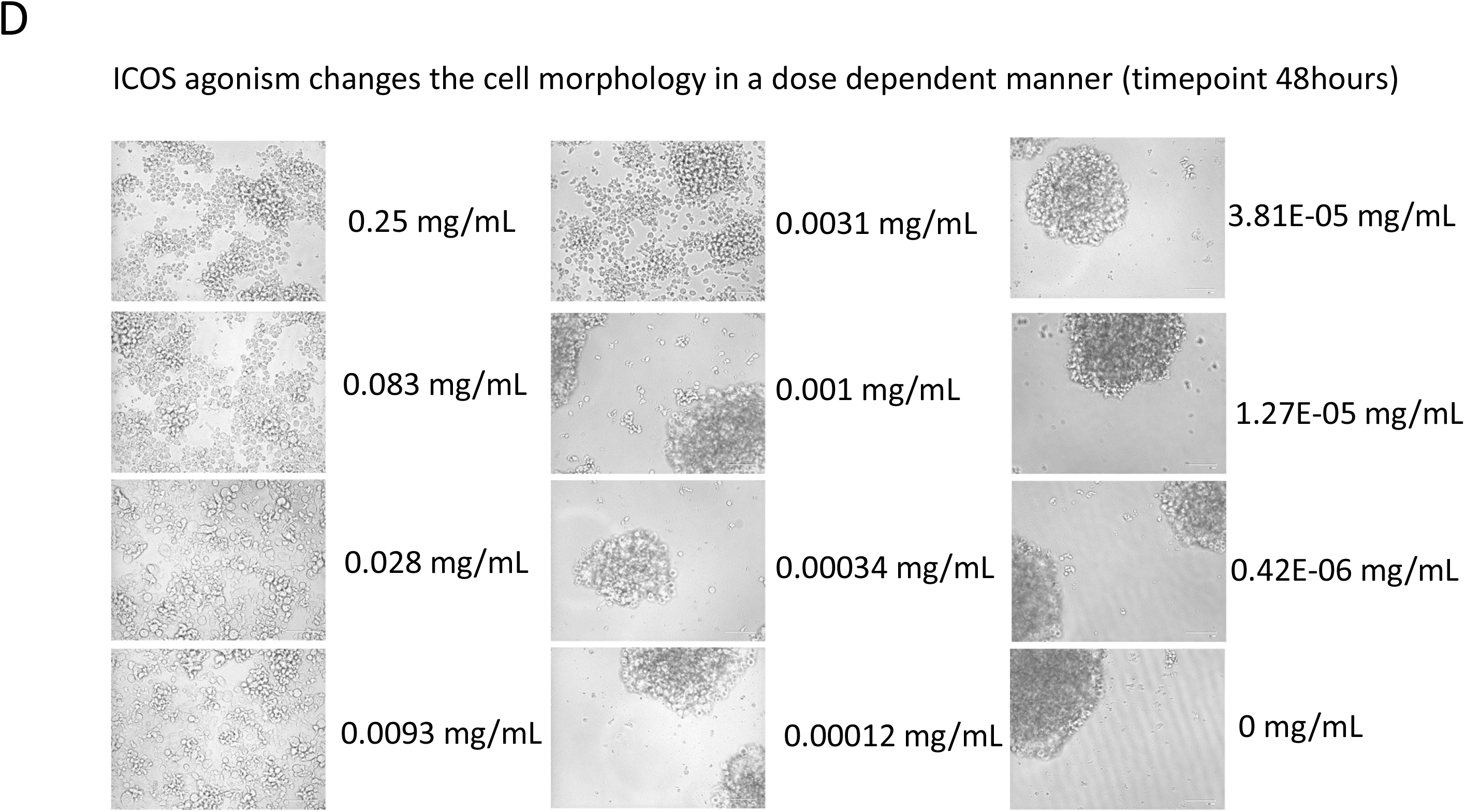
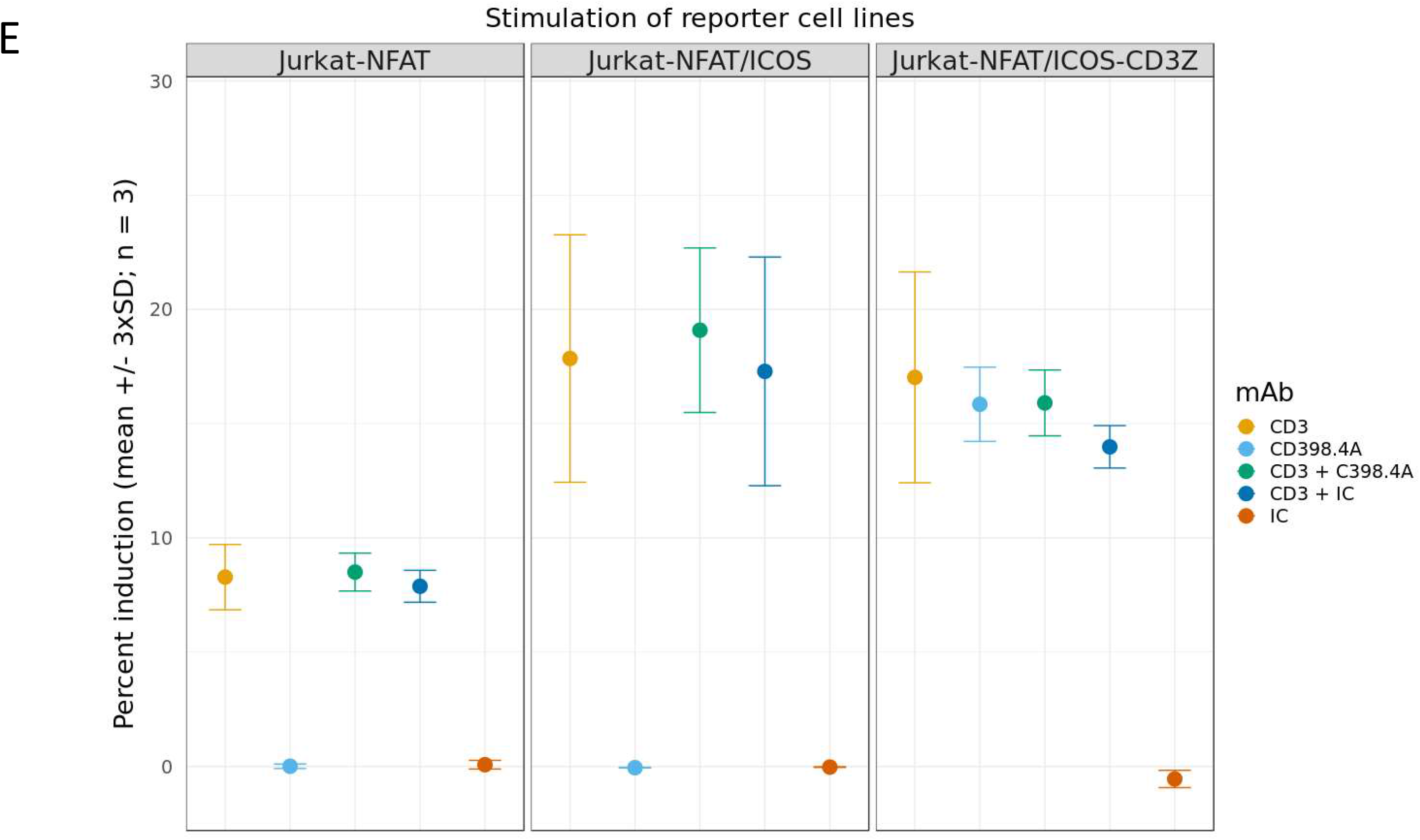
Percentage of ICOS positive cells and relative ICOS expression on CD4 and CD8 T cells at baseline (Day 0) and at day 3 and day 6 post anti-CD3/CD28 stimulation (A and B). (C) Time lapse microscopy over 24 hours of MJ cells cultured on plate precoated with an isotype control or with anti-ICOS C398.4A (20μg/ml). (D) images capture after 48 hours of MJ cells grown on plates pre-coated with different concentrations of KY1044. (E) Graph showing the NFAT luciferase reporter assays using untransfected or ICOS and ICOS-CD3ζ transfected Jurkat cells treated with different plate bound stimuli including anti-CD3 and anti-ICOS (C398.4A)

**Supplementary figure 3:**
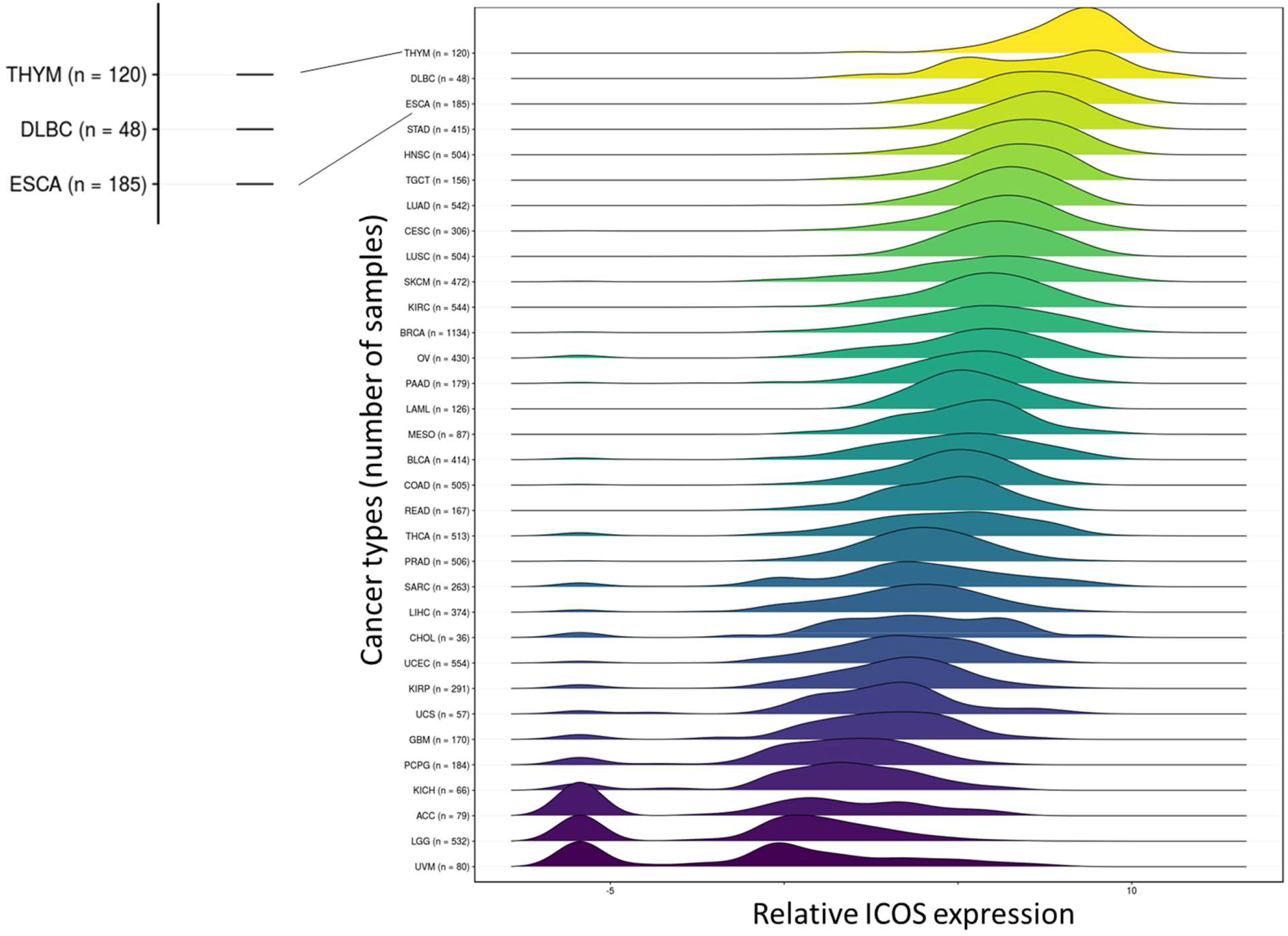
Density distributions showing the levels of *Icos* expression in haematological and solid tumour types from the TCGA datasets.

**Supplementary figure 4:**
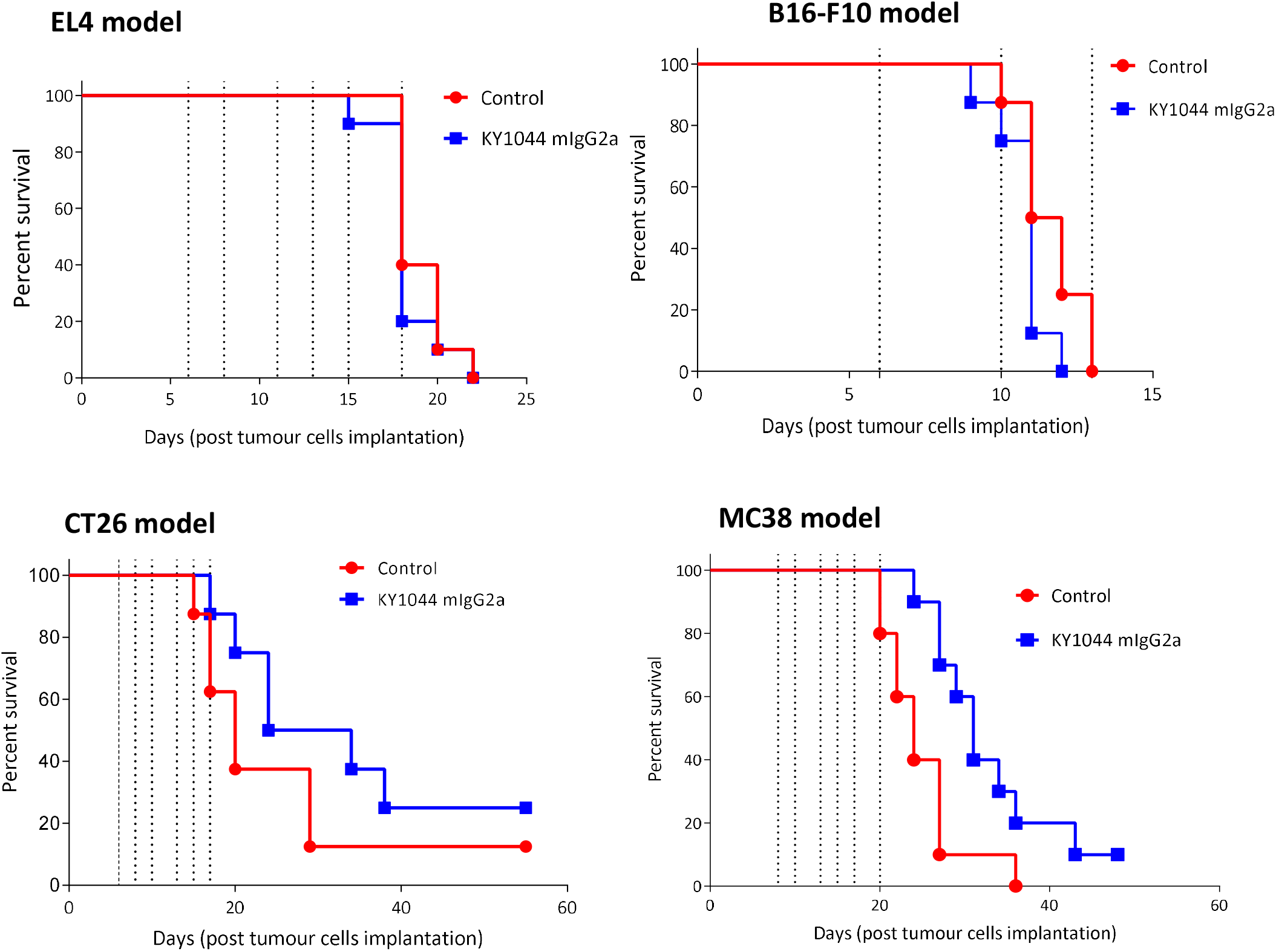
Examples of syngeneic tumour models resistant or poorly responsive to KY1044 mIgG2a monotherapy. Kaplan-Meier plot depicting the survival of mice injected sub-cut with different syngeneic tumour types (EL4, B16-F10, CT26 and MC38) and treated with KY1044 mIgG2a (3 or 10 mg/kg, vertical dotted lines).

**Supplementary figure 5:**
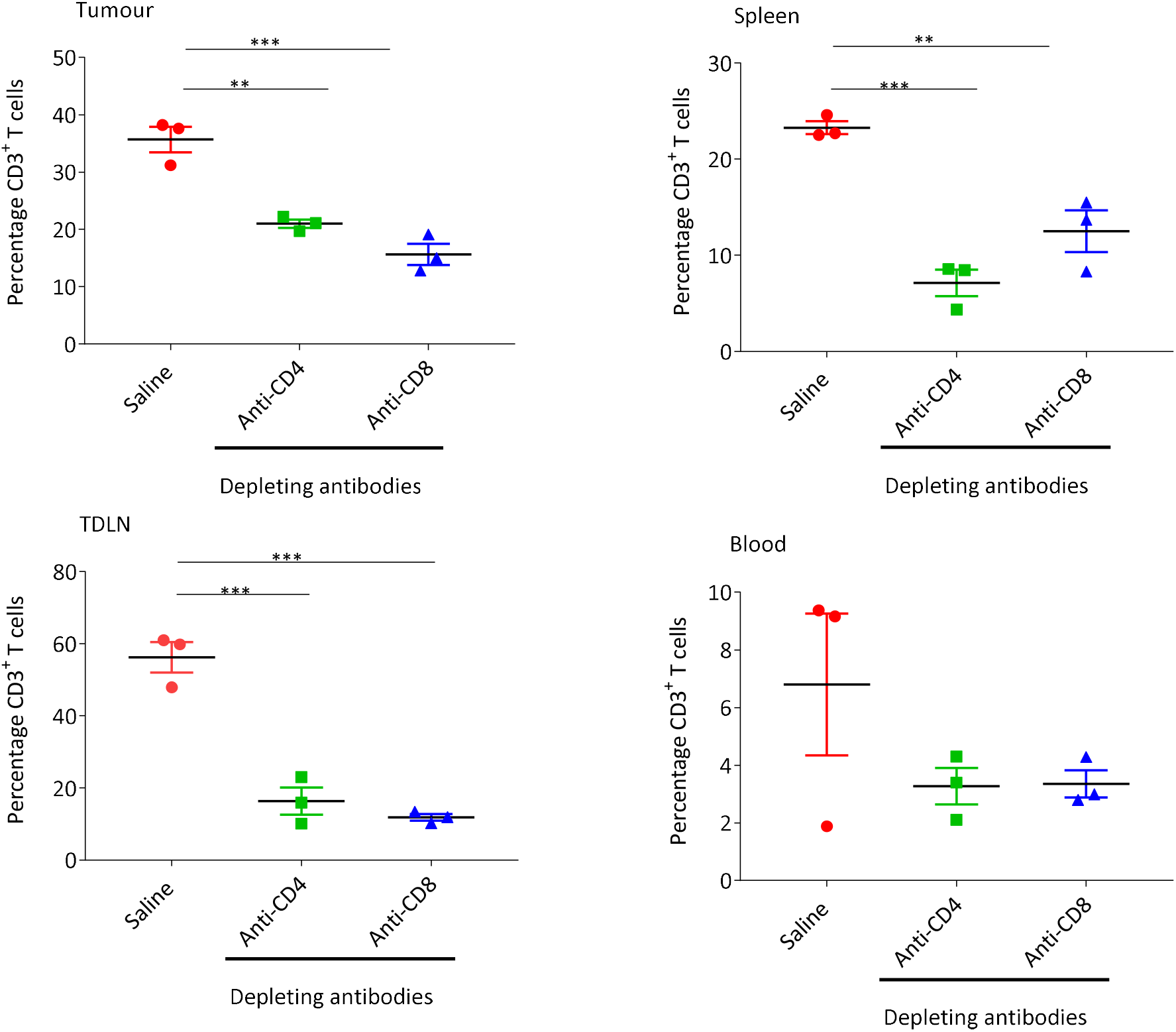
Graph summarising the analysis of T cells content in different mouse tissues (tumour, spleen, tumour draining lymph node [TDLN] and blood) treated with an isotype control antibody or anti-CD4 and anti-CD8 depleting antibodies. *** = P value ≤0.001 and ** = P value ≤0.01 (2-way ANOVA with Tukey’s multiple comparison).

**Supplementary figure 6:**
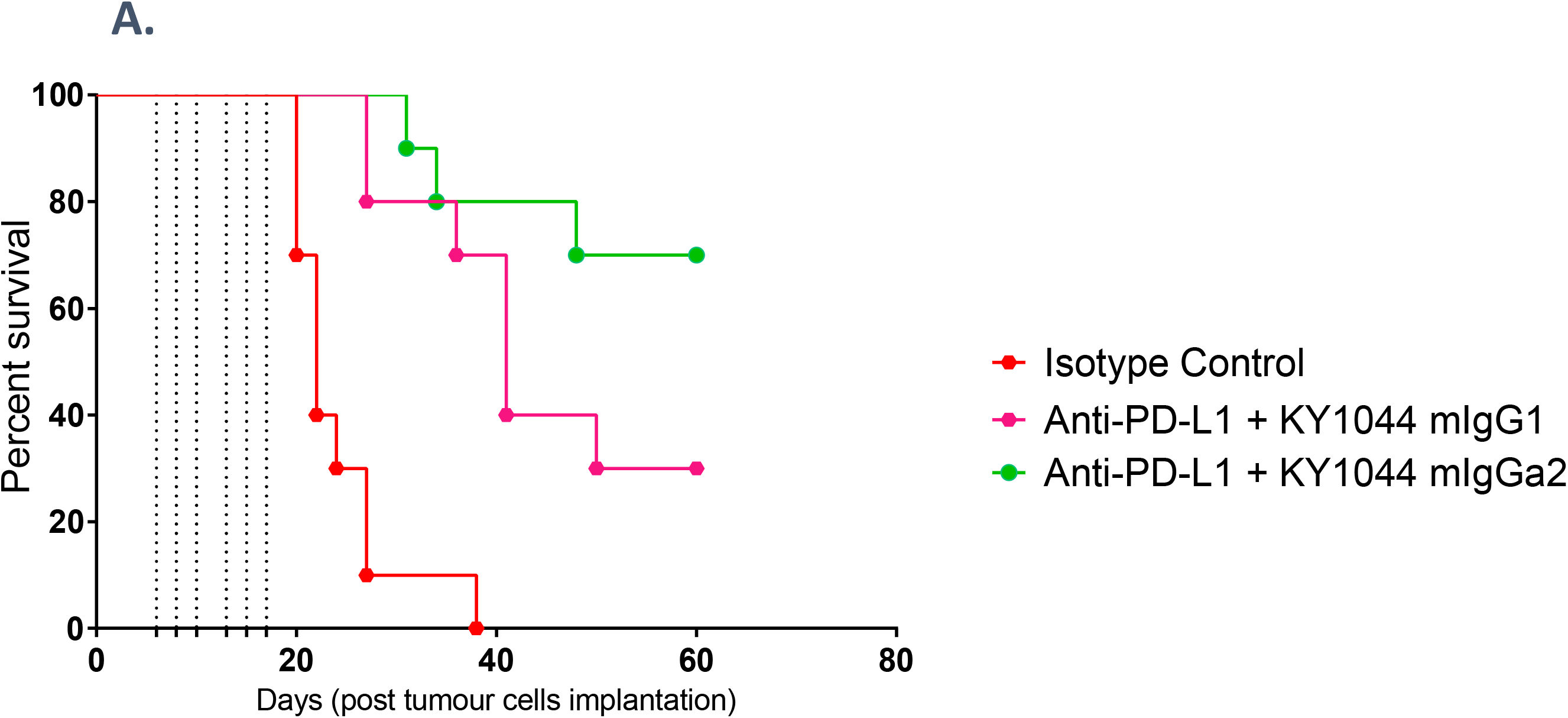
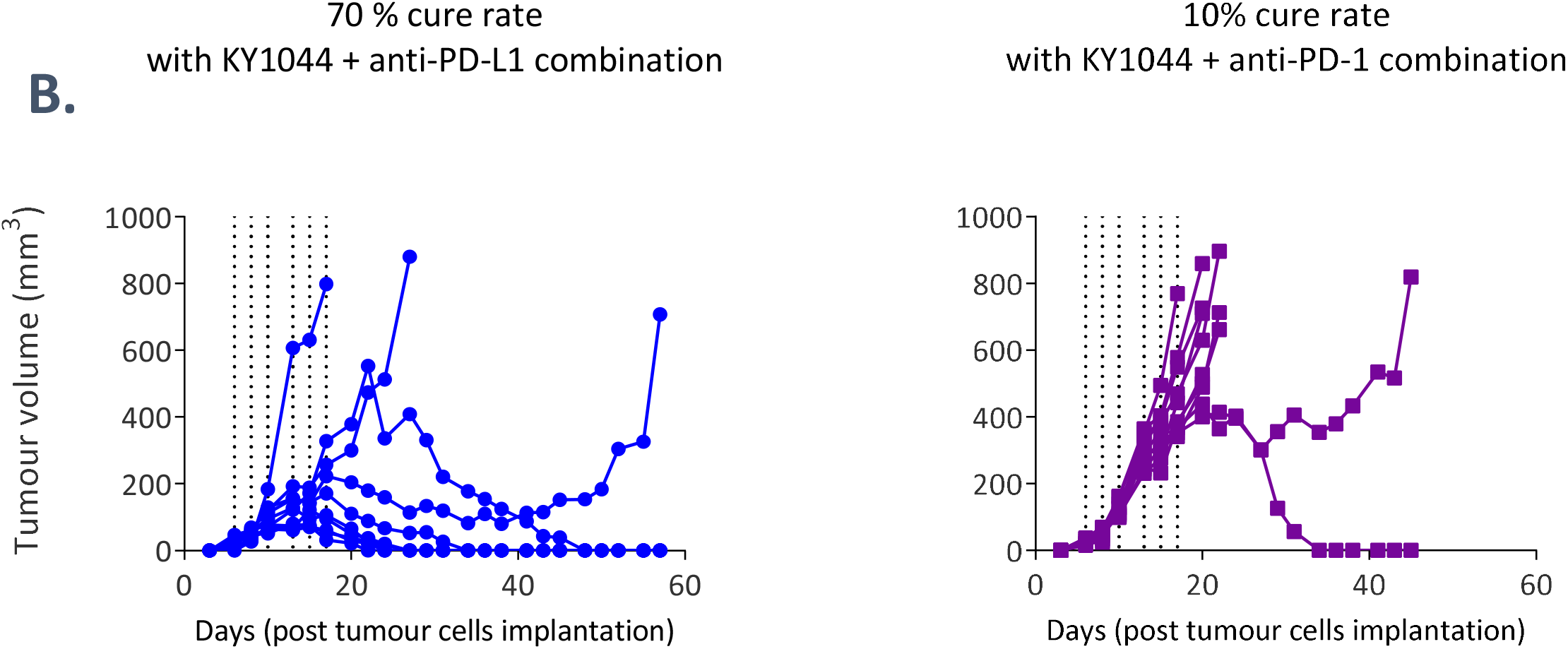
(A) Kaplan-Meier plot depicting the survival of mice injected sub-cut with CT26.WT syngeneic tumour cells line and treated with isotype control, KY1044 mIgG2a and anti-PD-L1 (3 and 10 mg/kg, respectively vertical dotted lines) or KY1044 mIgG1 and anti-PD-L1 (3 and 10 mg/kg, respectively vertical dotted lines). (B) spider plots showing the growth of CT26 tumours implanted in mice and treated with a combination of KY1044 mIgG2a with anti-PD-L1 (equivalent of 3 and 10 mg/kg, respectively vertical dotted lines, left) or a combination of KY1044 mIgG2a with anti-PD1(clone RMP1.14, equivalent of 3 and 10 mg/kg, respectively vertical dotted lines).

**Supplementary figure 7:**
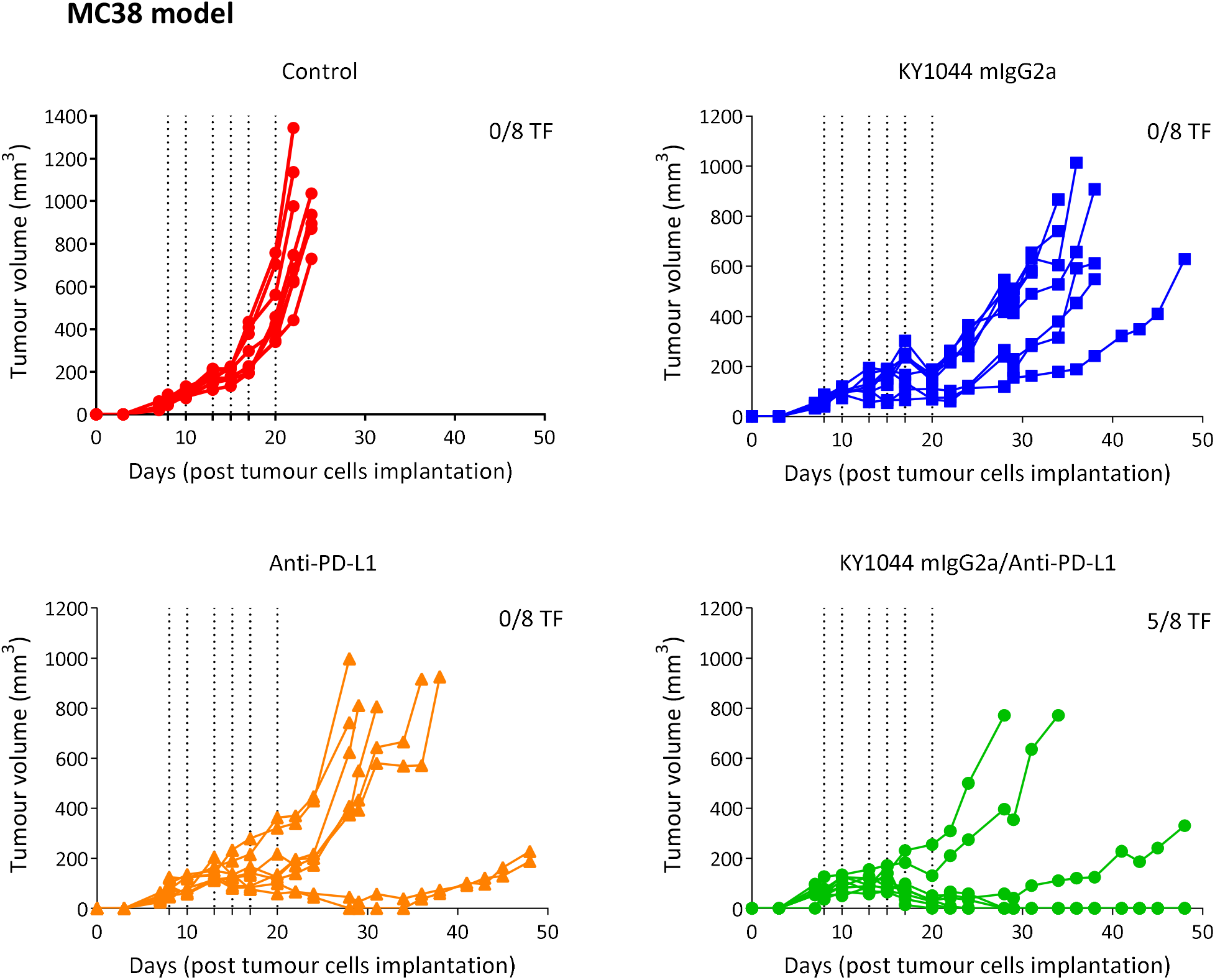
The combination of KY1044 mIgG2a with anti-PD-L1 is highly effective in the MC38 syngeneic tumour model. Spider plots showing individual mouse tumour volumes from C57Bl/6 mice (n=8 per group) harbouring subcutaneous MC38 tumours. Tumour bearing mice were dosed i.p (vertical dotted lines) with 60 µg of KY1044 mIgG2a and/or 200 µg of anti-PD-L1 starting from day 8 post tumour cell implantation

**Supplementary figure 8:**
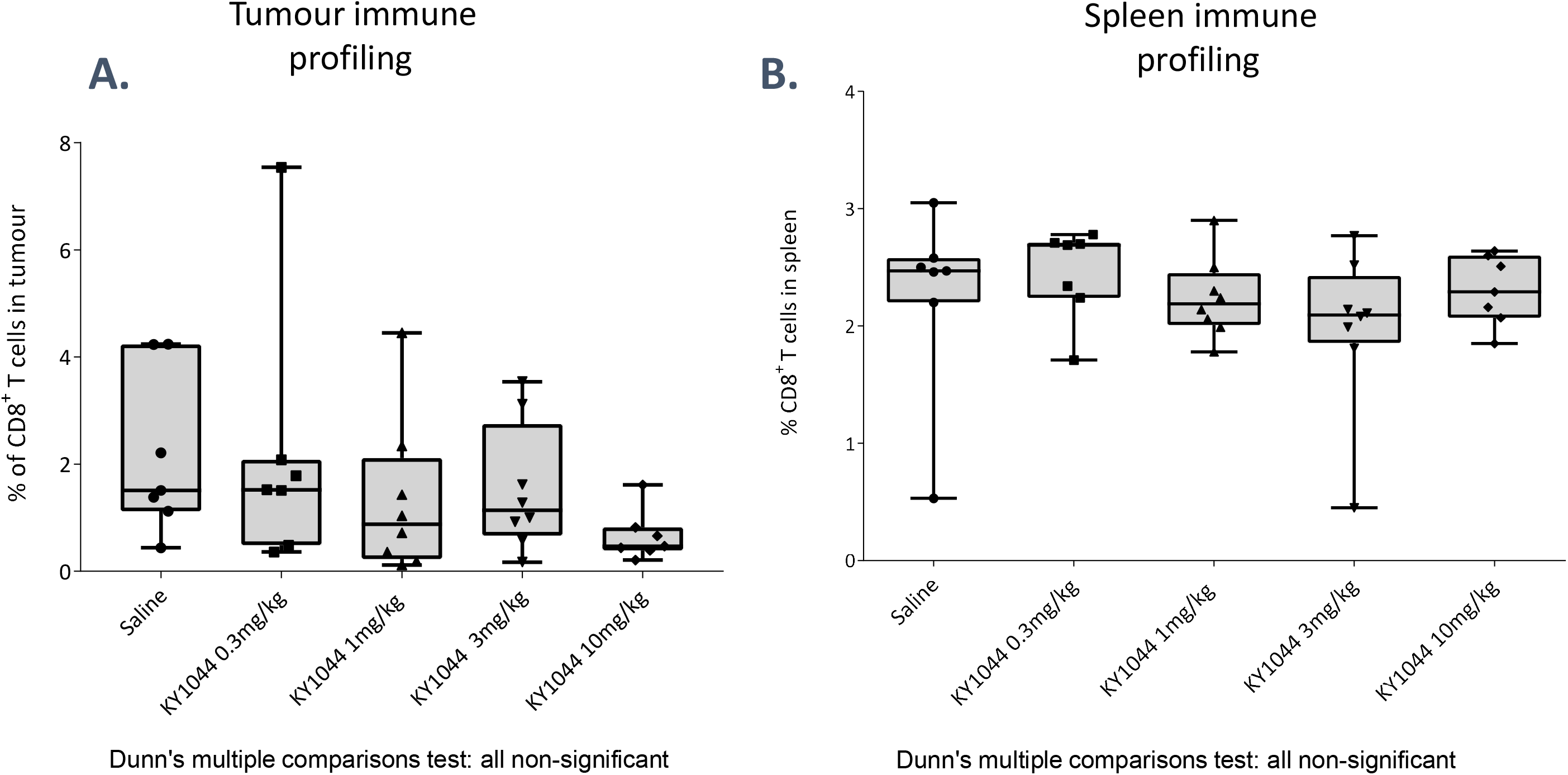
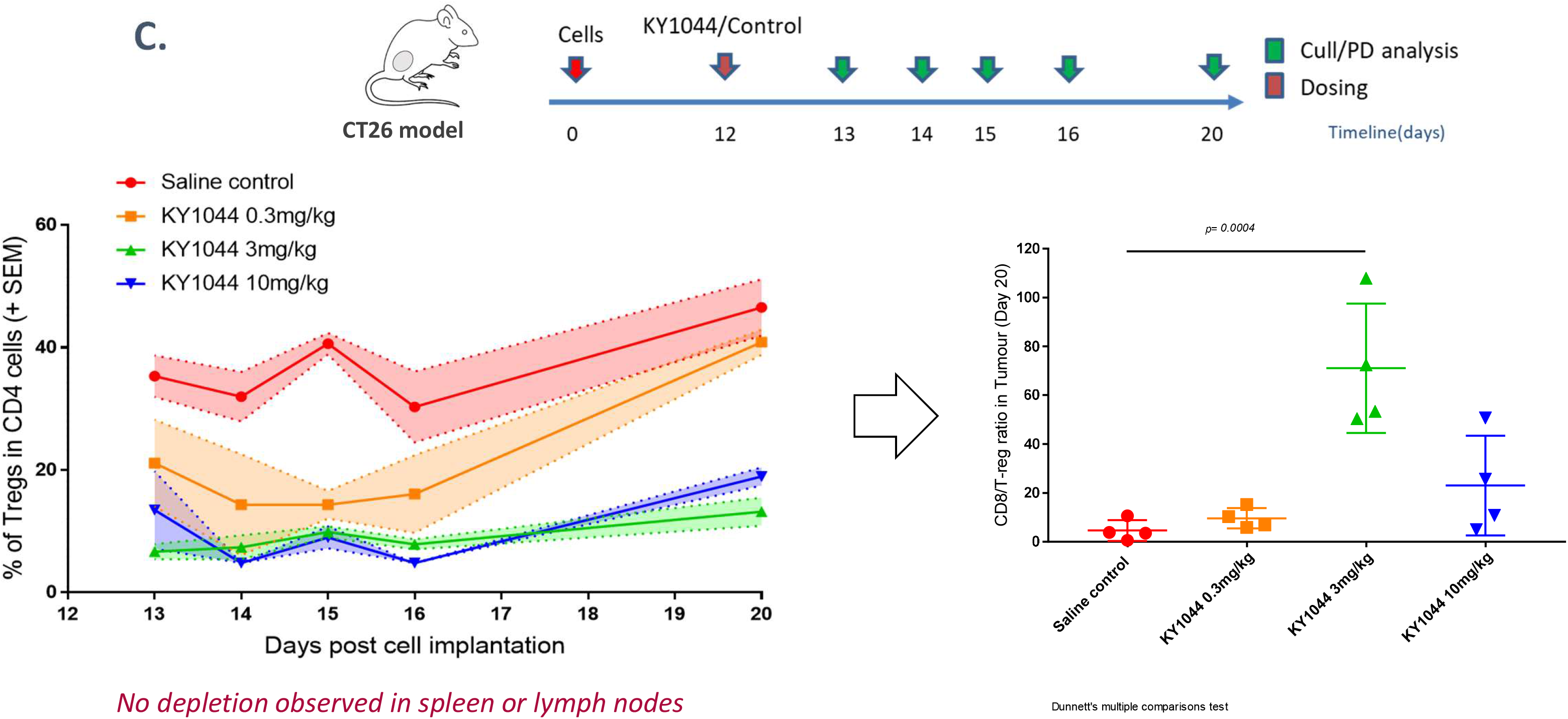
Flow cytometry analysis showing the percentage of CD8 T cells in the tumour and spleen of CT26 tumour bearing mice treated with different doses of KY1044. High dose KY1044 only marginally affects the percentage of CD8^+^ T in CT26 tumours but not in the spleen of tumour bearing mice (n=7/8 mice per groups) (A and B). (C) Longitudinal in vivo pharmacodynamic study looking at the T_regs_ content over time and the CD8 to T_reg_ ratio (Day 20) in CT26 tumours treated with different doses of KY1044. Note that the intermediate dose of 3mg/kg was associated with long term Treg depletion and the highest increase in the CD8 to T_reg_ ratio. (2-way ANOVA with Tukey’s multiple comparison).

**Supplementary figure 9:**
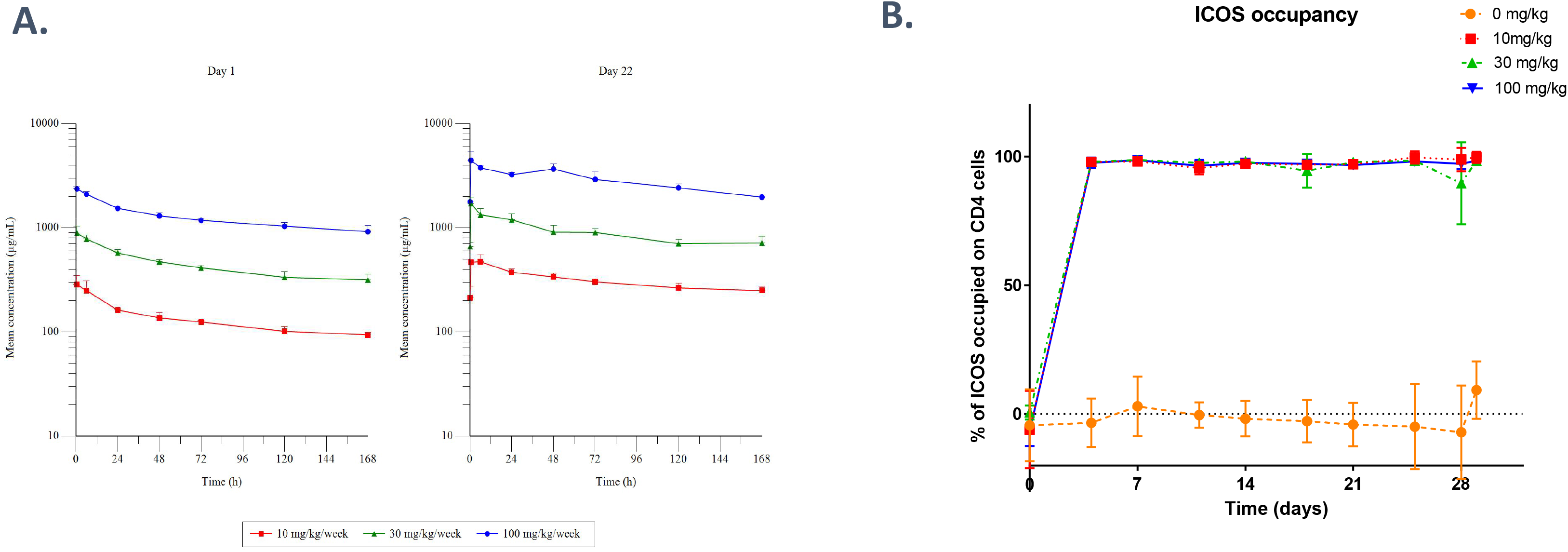
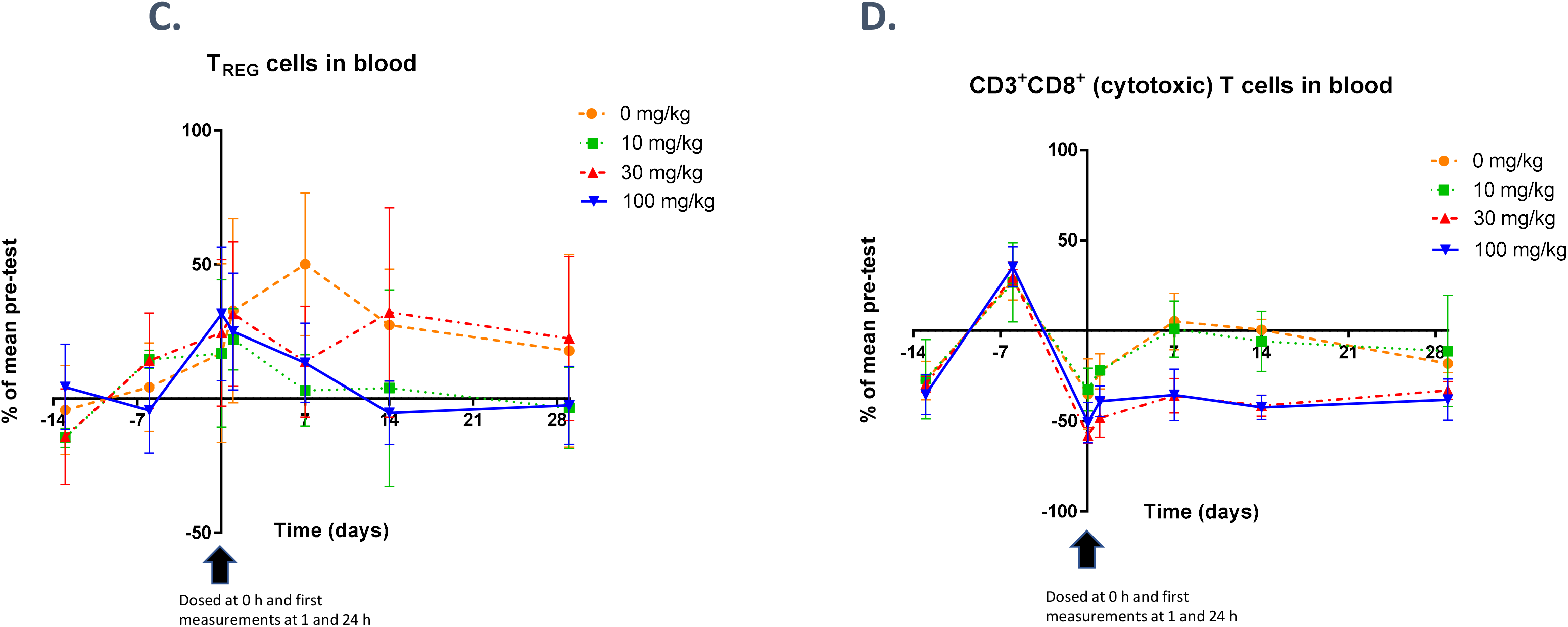
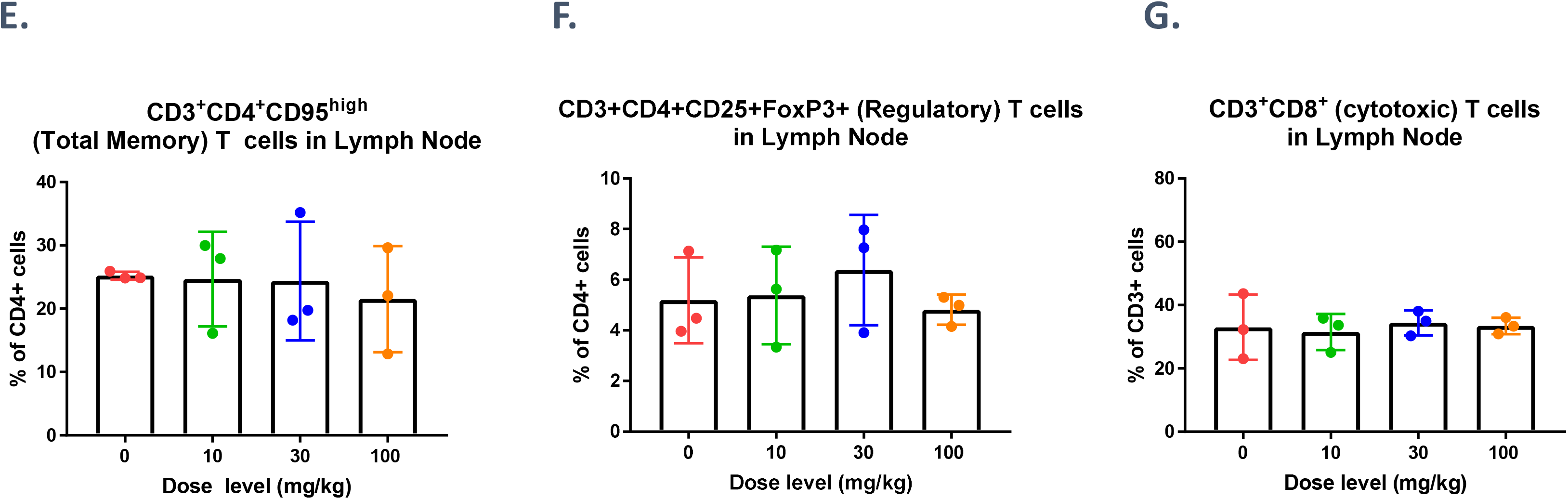
KY1044 hIgG1 exhibits dose-proportional linear pharmacokinetics in cynomolgus monkeys after weekly repeated i.v. dosing (serum profiles after the first and penultimate dose (Days 1 and 22; A) and full ICOS occupancy was maintained on blood CD4+ cells for all treated animals for the duration of the study (B). KY1044 hIgG1 does not have a notable effect on circulating T_Reg_ or CD8+ T cells in blood (C and D) or the proportion of T_M_, T_Reg_ or CD8+ T cells in the lymph nodes of NHP (E to G).

Supplementary table 1: Table summarizing the genes used to define the 27 immune cell subtypes analysed from the single cell RNA sequencing data generated from paired PBMC and tumour samples from 5 NSCLC cancer patients.

